# Comparison of the tolerability of ^161^Tb- and ^177^Lu-labeled somatostatin analogues in the preclinical setting

**DOI:** 10.1101/2024.03.29.587329

**Authors:** Sarah D. Busslinger, Ana Katrina Mapanao, Kristel Kegler, Peter Bernhardt, Fabienne Flühmann, Julia Fricke, Jan Rijn Zeevaart, Ulli Köster, Nicholas P. van der Meulen, Roger Schibli, Cristina Müller

## Abstract

**Purpose:** [^177^Lu]Lu-DOTATATE is an established somatostatin receptor (SSTR) agonist for the treatment of metastasized neuroendocrine neoplasms, while the SSTR antagonist [^177^Lu]Lu-DOTA-LM3 has only scarcely been employed in clinics. Impressive preclinical data obtained with [^161^Tb]Tb-DOTA-LM3 in tumor-bearing mice indicated the potential of terbium-161 as an alternative to lutetium-177. The aim of the present study was to compare the tolerability of ^161^Tb- and ^177^Lu-based DOTA-LM3 and DOTATATE in immunocompetent mice.

**Methods:** Dosimetry calculations were performed based on biodistribution data of the radiopeptides in immunocompetent mice. Treatment-related effects on blood cell counts were assessed on Days 10, 28 and 56 after application of [^161^Tb]Tb-DOTA-LM3 or [^161^Tb]Tb-DOTATATE at 20 MBq per mouse. These radiopeptides were also applied at 100 MBq per mouse and the effects compared to those observed after application of the ^177^Lu-labeled counterparts. Bone marrow smears, blood plasma parameters and organ histology were assessed at the end of the study.

**Results:** The absorbed organ dose was commonly higher for the SSTR antagonist than for the SSTR agonist and for terbium-161 over lutetium-177. Application of a therapeutic activity level of 20 MBq [^161^Tb]Tb-DOTA-LM3 or [^161^Tb]Tb-DOTATATE was well tolerated without major hematological changes. The injection of 100 MBq of the ^161^Tb- and ^177^Lu-based somatostatin analogues affected the blood cell counts, however. The lymphocytes were 40-50% lower in treated mice compared to the untreated controls on Day 10 irrespective of the radionuclide employed. At the same timepoint, thrombocyte and erythrocyte counts were 30–50% and 6–12% lower, respectively, after administration of the SSTR antagonist (*p*<0.05) while changes were less pronounced in mice injected with the SSTR agonist. All blood cell counts were in the normal range on Day 56. Histological analyses revealed minimal abnormalities in the kidneys, liver and spleen of treated mice. No correlation was observed between the organ dose and frequency of the occurrence of abnormalities.

**Conclusion:** Hematologic changes were more pronounced in mice treated with the SSTR antagonist than in those treated with the SSTR agonist. Despite the increased absorbed dose delivered by terbium-161 over lutetium-177, [^161^Tb]Tb-DOTA-LM3 and [^161^Tb]Tb-DOTATATE should be safe at activity levels that are recommended for their respective ^177^Lu-based analogues.

## Introduction

The expression of the somatostatin receptor (SSTR) in most neuroendocrine neoplasms (NENs) has been exploited for peptide receptor radionuclide therapy (PRRT) for more than two decades [1]. After initial application of [^111^In]In-DTPA-octreotide for Auger-electron therapy [2], the β^–^-particle-emitting yttrium-90 was employed for a more effective PRRT in combination with the SSTR agonists DOTATATE and DOTATOC [3]. The high-energy β^–^-particles emitted by yttrium-90 (Eβ^–^average = 932 keV) comprised the risk of radiation-induced nephropathy, however [4]. This concern was largely resolved by the introduction of lutetium-177 (Eβ^–^_average_ = 134 keV), which is currently the most often used radiometal for PRRT [5]. PRRT using ^177^Lu-based somatostatin analogues is commonly well tolerated and grade ≥3 adverse events have rarely been reported [6]. In most patients only mild hematological changes occurred with a recovery onset 4–6 weeks after radiopeptide application and cases of severe myelosuppression were the exception [6, 7].

SSTR antagonists, such as NODAGA/DOTA-LM3 and NODAGA/DOTA-JR11, have been developed and applied in combination with gallium-68 and lutetium-177 for imaging and treatment of NEN patients, respectively [8, 9]. Preclinically it was shown that, in contrast to SSTR agonists, SSTR antagonists bind to both the active and inactive forms of the SSTR and, hence, they show higher tumor uptake and better treatment efficacy [10, 11]. These findings were also confirmed in clinical studies with patients suffering from SSTR-positive malignancies [12]. Hematological changes were more often seen after application of SSTR antagonists than after standard PRRT using SSTR agonists, however [13, 14]. In order not to exceed a bone marrow dose of 1 Gy, the activity of ∼7.4 GBq per therapy cycle commonly applied with SSTR agonists had to be reduced to ∼4.5 GBq per therapy cycle when using SSTR antagonists [12, 14].

Complete response to PRRT using somatostatin analogues is rarely achieved [6], which may be ascribed to cancer cells that escape the treatment and develop to new metastases [15]. It is believed that this concern could - at least partially - be resolved by the application of terbium-161 instead of lutetium-177. Terbium-161 emits β^–^-particles as well as short-ranged electrons (conversion and Auger electrons) [15-18], which lead to an increased absorbed dose to single cancer cells and cancer cell clusters [15]. Preclinical therapy studies demonstrated the superiority of [^161^Tb]Tb-DOTATOC and [^161^Tb]Tb-DOTA-LM3 over their ^177^Lu-labeled counterparts and [^161^Tb]Tb-DOTA-LM3 was most effective to delay tumor growth in mice [19]. This situation motivated the translation of [^161^Tb]Tb-DOTA-LM3 to a clinical Phase 0/I study (NCT05359146) at the Basel University Hospital in Switzerland. Initial imaging data showed promising results in terms of tumor uptake and retention of [^161^Tb]Tb-DOTA-LM3 [20].

The question of whether terbium-161 enhances the risk of bone marrow toxicity remains unexplored. It was, therefore, our goal to investigate treatment-induced adverse events in immunocompetent mice after application of 20 MBq [^161^Tb]Tb-DOTA-LM3 or 20 MBq [^161^Tb]Tb-DOTATATE, an activity that would − according to a body mass-based and surface-related extrapolation [21] − translate to ∼4.5 GBq for a human patient. This activity corresponds well with the activity of a ^177^Lu-based SSTR antagonist (∼4.5 GBq/cycle) in recently published clinical studies (NCT02592707 and NCT04997317) [12, 14]. The ^161^Tb-based radiopeptides were also investigated at a 5-fold higher activity (100 MBq per mouse) to compare hematological changes with those observed after application of the ^177^Lu-labeled counterparts. Potential damage to the kidneys and other normal organs and tissues was also assessed.

## Materials and methods

### Radiopeptides

Terbium-161 (produced in-house as previously reported [22] or obtained from Terthera, the Netherlands, via Solumedics AG, Switzerland) and lutetium-177 (ITM Medical Isotopes GmbH, Germany) were measured using calibrated instrumentation (Supplementary Material) [23]. DOTA-LM3 (CSBio, Silicon Valley Menlo Park, California, United States) and DOTATATE (Bachem, Bubendorf, Switzerland) were labeled with terbium-161 or lutetium-177 at a molar activity up to 100 MBq/nmol using standard labeling conditions as previously reported (Supplementary Material) [19].

### In vivo studies

All applicable international, national, and/or institutional guidelines for the care and use of laboratory animals, in particular the guidelines of Swiss Regulations for Animal Welfare were applied. The preclinical studies were ethically approved by the responsible Committee of Animal Experimentation and permitted by the responsible cantonal authorities (license N° 75721 and N° 79692). Five-to-six-week-old immunocompetent, female FVB mice were purchased from Charles River Laboratories (Sulzfeld, Germany) and acclimatized for at least one week after arrival at the Paul Scherrer Institute.

### Biodistribution study

Immunocompetent FVB mice were intravenously injected with [^161^Tb]Tb-DOTA-LM3 (5 MBq, 1 nmol) or [^161^Tb]Tb-DOTATATE (5 MBq, 1 nmol) diluted in phosphate-buffered saline (PBS pH 7.4) containing 0.05% bovine serum albumin. The mice were dissected at 2 h, 6 h, 24 h or 72 h after injection of the radiopeptides. Collected tissues and organs were weighed and counted for activity using a γ-counter (PerkinElmer, Wallac Wizard 1480, Waltham, MA, United States). Decay-corrected values of the injected activity per gram tissue mass (% IA/g) were calculated as the average ± standard deviation (SD) of n=3 mice per timepoint (Supplementary Material).

### In vitro autoradiography of bone marrow cells

Bone marrow cells were isolated from the femurs, tibiae and iliac bones of immunocompetent FVB mice according to a previously published protocol (Supplementary Material) [24]. The bone marrow cells (5 × 10^6^ cells per sample) were incubated with the radiopeptides (50 kBq, 0.5 pmol) in a volume of 1 mL in the absence or presence of excess unlabeled peptide for 2 h at 37 °C. After rinsing with PBS, the bone marrow cells were placed on a membrane and left to dry. Images were obtained by exposure of the cells to a super resolution phosphor imaging screen (SR 7001486, PerkinElmer, Waltham, MA, United States) for 72 h, followed by screen reading using a storage phosphor system (Cyclone Plus, PerkinElmer, Waltham, MA, United States). The detected activity, corresponding to the cell-bound radiopeptide fraction, was quantified using OptiQuant software (version 5.0, Bright Instrument Co Ltd., PerkinElmer, Waltham, MA, United States). The data were presented as average ± SD of 3 independent experiments, each performed in duplicate and in parallel for both radiopeptides using bone marrow cells of one mouse per experiment. The experimental setting was validated using SSTR-negative KB tumor cells mixed with a small fraction of SSTR-positive AR42J tumor cells (Supplementary Material), in order to simulate the situation in the bone marrow which is reported to contain a small fraction of SSTR-positive cells [25].

### Dosimetry calculations

The absorbed dose in relevant organs was calculated based on non-decay-corrected biodistribution data obtained for [^161^Tb]Tb-DOTA-LM3 and [^161^Tb]Tb-DOTATATE. The accumulated activity was plotted against the time and the bi-exponential curve fits were integrated to infinity to obtain the time-integrated activity concentration (TIAC). The absorbed energy per decay was obtained from the mouse phantom MOBY [26]. The activity concentration in the bone marrow was calculated by using the mass relation between the cortical bone and bone marrow in the femur in the mouse phantom, assuming equal activity concentration in the cortical bone and muscle. As the tissue distribution of the radiopeptide remained the same, irrespective of the employed radionuclide [27], the absorbed doses for [^177^Lu]Lu-DOTA-LM3 and [^177^Lu]Lu-DOTATATE were calculated by taking the difference in absorbed energy and physical half-life between the two radionuclides into account (Supplementary Material).

### Design of the tolerability study in mice

In Study I, the tolerability of [^161^Tb]Tb-DOTA-LM3 and [^161^Tb]Tb-DOTATATE was tested in immunocompetent FVB mice (n=7 per group) at an activity of 20 MBq (0.2 nmol) per mouse (Supplementary Material). In Study II, additional mice (n=7 per group) were injected with [^161^Tb]Tb-DOTA-LM3 and [^161^Tb]Tb-DOTATATE and the ^177^Lu-labeled analogues, applied at 5-fold higher activity (100 MBq; 1 nmol per mouse). In both studies, mice were monitored over 56 days after application of the radiopeptides.

### Monitoring of mice during Study I and Study II

The mice were monitored throughout the entire study regarding the body mass, behavior and signs of unease or pain using a scoring system to assess whether pre-defined endpoint criteria were reached. The body mass was measured three times a week and expressed relative to the body mass determined on the day of injection (Day 0; set as 1.0; Supplementary Material). At study end on Day 56, the brain, spleen, kidneys and liver were collected and weighed to calculate the organ-to-brain mass ratios.

Blood cell counts were determined on Days 10, 28 and 56 after treatment using a hematology analyzer (VetScan HM5, Abaxis, United States) and corresponding blood smears were prepared. Bone marrow smears were prepared after euthanasia on Day 56. The smears were stained using the Pappenheim method [28]. In Study II, potential morphological changes of the blood cells were assessed by a board-certified veterinary pathologist (AnaPath Services GmbH, Liestal, Switzerland; Supplementary Material). Blood plasma parameters were determined at study end using a dry chemistry analyzer (DRI-CHEM 4000i, FUJIFILM, Japan) as previously reported (Supplementary Material) [19].

Tissue sections (4 µm thickness) of paraffin-embedded spleen, kidneys and liver of mice from Study II were stained using hematoxylin and eosin and the histo(patho)logical analysis was performed by the veterinary pathologist using a pre-defined scoring system (Supplementary Material) [29].

### Statistical analysis

The blood cell counts were analyzed using a two-way ANOVA, whereas for all other data, a one-way ANOVA was applied. A Tukey’s multiple comparisons post-test was used for comparison of the individual groups (GraphPad Prism software, version 7.0). A *p* value ≤0.05 was considered statistically significant.

## Results

### Radiopeptides

The radiochemical purity of the radiopeptides employed herein was >98% at a molar activity of up to 100 MBq/nmol, irrespective of whether terbium-161 or lutetium-177 was used (Supplementary Material, Fig. S1). This allowed the use of the radiopeptides without further purification for preclinical investigations.

### Biodistribution study

Rapid blood clearance of [^161^Tb]Tb-DOTA-LM3 and [^161^Tb]Tb-DOTATATE resulted in blood values of <0.2% IA/g at 2 h post injection (p.i.). Both radiopeptides showed high renal retention, with values about twice as high for [^161^Tb]Tb-DOTA-LM3 (17 ± 2% IA/g at 2 h p.i. and 8.3 ± 0.6% IA/g at 24 h p.i.) than for [^161^Tb]Tb-DOTATATE (9.0 ± 0.6% IA/g at 2 h p.i. and 3.4 ± 0.5% IA/g at 24 h p.i.). In the spleen and bone marrow as part of the femur, [^161^Tb]Tb-DOTA-LM3 showed higher accumulation than [^161^Tb]Tb-DOTATATE. On the other hand, the uptake of the two radiopeptides was similar in the adrenals and in the thymus. In the lungs and liver, the retention of [^161^Tb]Tb-DOTA-LM3 was higher than that of [^161^Tb]Tb-DOTATATE. Aside from the kidneys, accumulution in other organs and tissues was <2% IA/g at 24 h after injection of either radiopeptide (Fig. 1, Supplementary Material, Tables S1 and S2).

**Fig. 1.**
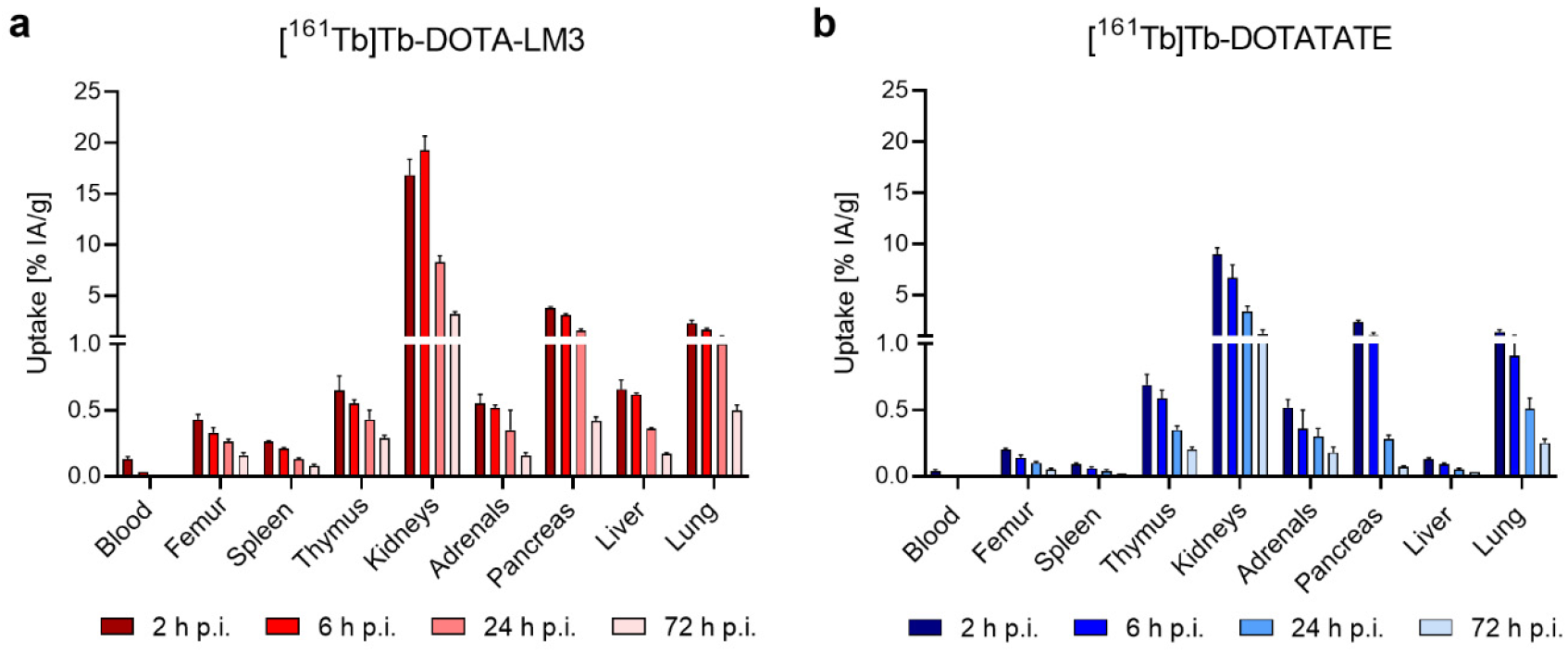
**a/b** Decay-corrected biodistribution data of the radiopeptides in immunocompetent FVB mice, shown as the average ± SD of n=3 mice; **a** Biodistribution data of [^161^Tb]Tb-DOTA-LM3; **b** Biodistribution data of [^161^Tb]Tb-DOTATATE.

### SSTR-specific binding of the radiopeptides to bone marrow cells

The SSTR-specific binding of [^177^Lu]Lu-DOTA-LM3 and [^177^Lu]Lu-DOTATATE to bone marrow cells was visualized using the technique of autoradiography (Supplementary Material, Fig. S2). The signal intensity correlated negatively with the amount of added unlabeled peptide to the cells, which blocked the SSTRs (Fig. 2, Supplementary Material, Fig. S3). A 10-fold excess of the respective unlabeled peptide reduced the binding of [^177^Lu]Lu-DOTA-LM3 and [^177^Lu]Lu-DOTATATE by 82% and 79%, respectively. Simulation experiments performed with SSTR-negative KB tumor cells mixed with a small fraction of SSTR-positive AR42J tumor cells showed a similar autoradiography pattern with a lower signal after coincubation of the cells with unlabeled peptide. The signal intensity, however, was considerably lower for bone marrow cells than for tumor cells indicating an extremely low expression level of the SSTR in bone marrow cells (Supplementary Material, Fig. S3).

**Fig. 2.**
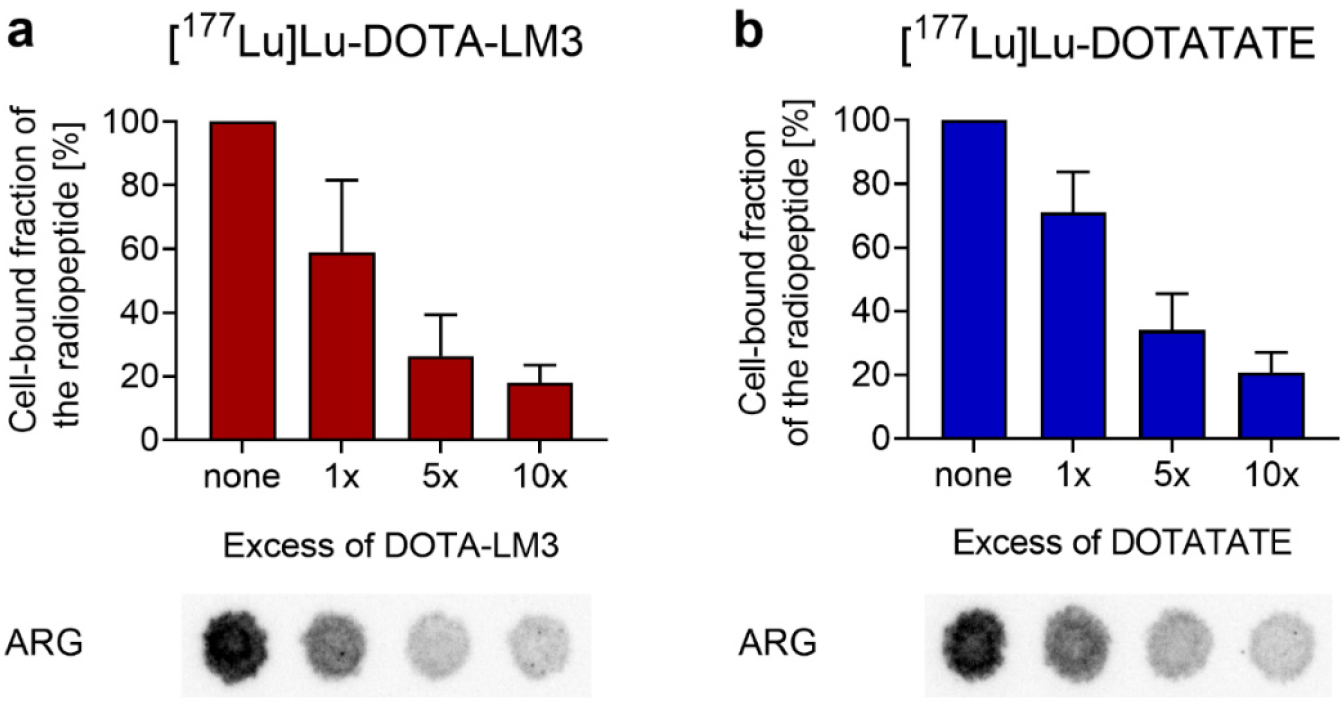
**a/b** Binding of the radiopeptides to bone marrow cells in the presence of excess unlabeled peptide. **a** [^177^Lu]Lu-DOTA-LM3; **b** [^177^Lu]Lu-DOTATATE. The binding of the radiopeptides is shown relative to the measurements of samples in the absence of unlabeled peptide (set as 100%). The results are presented as average ± SD of 3 independent experiments. Representative autoradiograms (ARG) of the respective samples are shown in the lower panel.

### Dosimetry estimations

Dosimetry calculations for [^161^Tb]Tb-DOTA-LM3 revealed a 3.2- to 3.6-fold higher absorbed dose to the SSTR-positive spleen, kidneys, pancreas and bone marrow (as part of the femur) than for [^161^Tb]Tb-DOTATATE. The absorbed dose to the thymus was similar for both radiopeptides. The absorbed dose to the lungs was 1.9-fold higher for the SSTR antagonist than for the SSTR agonist and the absorbed dose to the liver was 8.6-fold higher for the SSTR antagonist than for the SSTR-agonist (Table 1). Relative to the absorbed dose of the ^161^Tb-based radiopeptides, the absorbed dose to the same organs was about 1.4-fold lower when using the ^177^Lu-labeled counterparts.

**Table 1.** Absorbed organ doses calculated based on non-decay-corrected biodistribution data obtained for [^161^Tb]Tb-DOTA-LM3 and [^161^Tb]Tb-DOTATATE and extrapolated for the ^177^Lu-labeled counterparts.

| Organ | [<sup>161</sup>Tb]Tb-DOTA-LM3 | [<sup>177</sup>Lu]Lu-DOTA-LM3 | [<sup>161</sup>Tb]Tb-DOTATATE | [<sup>177</sup>Lu]Lu-DOTATATE | Dose ratio DOTA-LM3/DOTATATE |
| --- | --- | --- | --- | --- | --- |
|  | [mGy/MBq] | [mGy/MBq] | [mGy/MBq] | [mGy/MBq] |  |
| Femur<sup>a</sup> | 47 ± 4 | 34 ± 3 | 15 ± 1 | 11 ± 1 | 3.2 |
| Spleen | 14 ± 1 | 10 ± 1 | 5 ± 1 | 3 ± 1 | 3.2 |
| Thymus | 49 ± 11 | 36 ± 8 | 51 ± 4 | 37 ± 3 | 1.0 |
| Kidneys | 1000 ± 100 | 760 ± 60 | 300 ± 40 | 220 ± 30 | 3.5 |
| Pancreas | 120 ± 10 | 84 ± 3 | 33 ± 1 | 24 ± 1 | 3.6 |
| Liver | 49 ± 5 | 36 ± 4 | 6 ± 1 | 4 ± 1 | 8.6 |
| Lung | 68 ± 3 | 50 ± 2 | 36 ± 3 | 26 ± 2 | 1.9 |

<sup>a</sup>Femur containing the bone marrow.

### Study I: Tolerability of [^161^Tb]Tb-DOTA-LM3 and [^161^Tb]Tb-DOTATATE (20 MBq/mouse)

In an initial study, the tolerability of [^161^Tb]Tb-DOTA-LM3 and [^161^Tb]Tb-DOTATATE at an activity of 20 MBq per mouse was investigated. According to body mass and surface-based extrapolation [21], this activity would correspond to ∼4.5 GBq radiopeptide for a human patient (Supplementary Material). The average body mass of mice treated with either [^161^Tb]Tb-DOTA-LM3 or [^161^Tb]Tb-DOTATATE increased over the entire study like for untreated controls (*p*>0.05; Supplementary Material, Fig. S4). At the time of euthanasia, the average brain, spleen and kidney masses and related organ-to-brain mass ratios of treated mice were in the same range as for untreated control mice, irrespective of the radiopeptide applied (*p*>0.05; Supplementary Material, Tables S3 and S4). Liver masses were slightly higher for treated mice than for untreated controls, which resulted in a significantly higher liver-to-brain mass ratio for [^161^Tb]Tb-DOTATATE in comparison to controls (*p*<0.05).

Blood cell counts were in the same range for mice treated with [^161^Tb]Tb-DOTA-LM3 and [^161^Tb]Tb-DOTATATE (*p*>0.05), but differences were seen in comparison to control values. On Day 10 after application of either radiopeptide, thrombocyte counts were lower ((393 ± 58) × 10^9^ cells/L (*p*<0.05) and (403 ± 92) × 10^9^ cells/L (*p*>0.05), respectively) than in control mice ((469 ± 52) × 10^9^ cells/L), but had fully recovered by Day 28. The erythrocyte counts were also somewhat lower in treated mice compared to untreated controls at Day 10 and Day 28 (*p*>0.05) but reached normal levels by Day 56. The same trend was seen for hemoglobin values with a significant drop for mice injected with [^161^Tb]Tb-DOTATATE (*p*<0.05). While average leukocyte counts were not significantly different between control mice and treated mice on Day 10, significantly fewer leukocytes were seen in treated mice ((7.0 − 7.9) × 10^9^ cells/L) than in untreated controls ((10.7 ± 1.3) × 10^9^ cells/L; *p*<0.05) on Day 28. Full recovery had occurred by Day 56, however. This transient decrease in leukocytes was ascribed to reduced lymphocytes (Fig. 3; Supplementary Material, Tables S5 and S6).

**Fig. 3.**
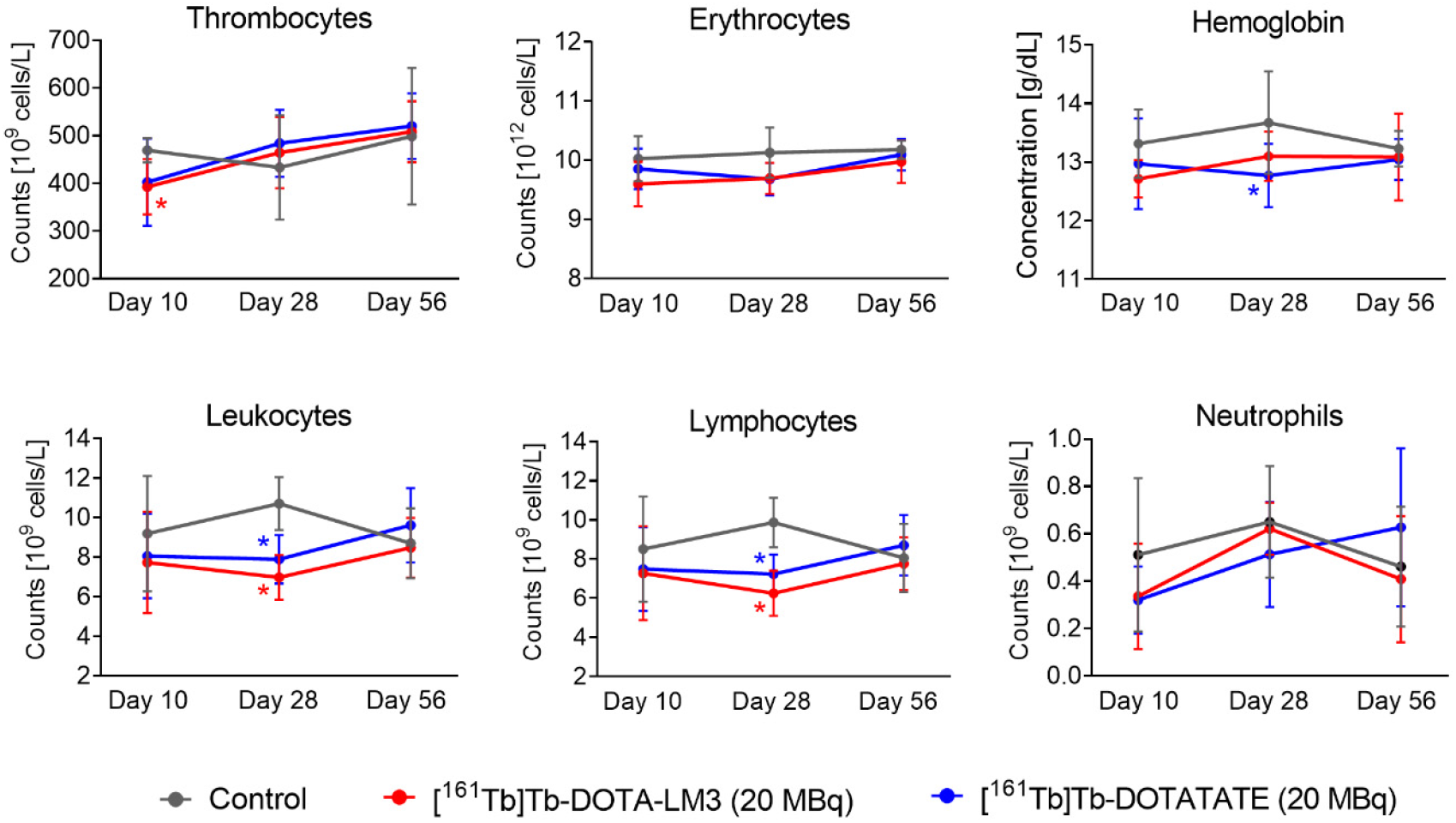
Blood cell counts and hemoglobin values on Day 10, Day 28 and Day 56 after treatment of mice with [^161^Tb]Tb-DOTA-LM3 or [^161^Tb]Tb-DOTATATE (20 MBq; 0.2 nmol per mouse). (*) Values significantly different from those of the control group.

Similar concentrations of blood urea nitrogen, alkaline phosphatase and albumin in blood plasma were determined for treated mice and untreated controls on Day 56 (*p*>0.05; Supplementary Material, Table S7).

### Study II: Tolerability of ^161^Tb- and ^177^Lu-based radiopeptides (100 MBq/mouse)

Potential differences in the tolerability of [^161^Tb]Tb-DOTA-LM3 and [^161^Tb]Tb-DOTATATE as compared to [^177^Lu]Lu-DOTA-LM3 and [^177^Lu]Lu-DOTATATE, respectively, were estimated in a follow-up study employing a 5-fold higher activity than in Study I. This activity would translate to approximately 22 GBq for a human patient [21], which corresponds to the cumulative activity of about 3 therapy cycles of [^177^Lu]Lu-DOTATATE (∼7.4 GBq/cycle) [6] and was, thus, much more than would ever be used for a single patient injection.

Overall, mice gained body mass over time and only one mouse that received [^161^Tb]Tb-DOTA-LM3 and one that received [^177^Lu]Lu-DOTATATE transiently lost 5-12% of the body mass, with lowest values reached about 2 weeks after injection (Supplementary Material, Fig. S5). Organ masses and organ-to-brain mass ratios at study end remained unaffected (*p*>0.05). Only the liver mass was enhanced in treated mice, which resulted in up to 27% higher liver-to-brain mass ratios than in untreated control mice (Supplementary Material, Tables S8 and S9).

Treated mice showed significantly lower thrombocyte counts compared to control mice (*p*<0.05) on Day 10 after injection (Fig. 4, Supplementary Material, Table S10). This was particularly the case for mice that received [^161^Tb]Tb-DOTA-LM3 or [^177^Lu]Lu-DOTA-LM3, for which the average values were 49% and 30% lower than that for mice of the control group, respectively. On Day 28, the thrombocyte counts of treated mice were, however, similar to those of untreated controls (*p*>0.05). Erythrocyte counts on Day 10 were not significantly different between mice treated with [^161^Tb]Tb-DOTA-LM3 and [^177^Lu]Lu-DOTA-LM3 (*p*>0.05), but these values were significantly lower than those of control mice (*p*<0.05). Values in the normal range were reached at study end, with the average cell counts being about 5% lower in treated mice than in untreated control mice (*p*<0.05). The erythrocyte counts of mice injected with [^161^Tb]Tb-DOTATATE or [^177^Lu]Lu-DOTATATE fall within the range measured for untreated controls at all investigated timepoints (*p*>0.05; Fig. 4; Supplementary Material, Table S10). A significant decrease in the hemoglobin concentration as compared to that of control mice was seen solely for mice treated with [^161^Tb]Tb-DOTA-LM3 (*p*<0.05) on Day 10 (Fig. 4). At this timepoint, all treated mice had lower hematocrit values than control mice (Fig. 4), but normal values were determined on Day 28 and on Day 56 (*p*>0.05; Supplementary Material, Table S10).

**Fig. 4.**
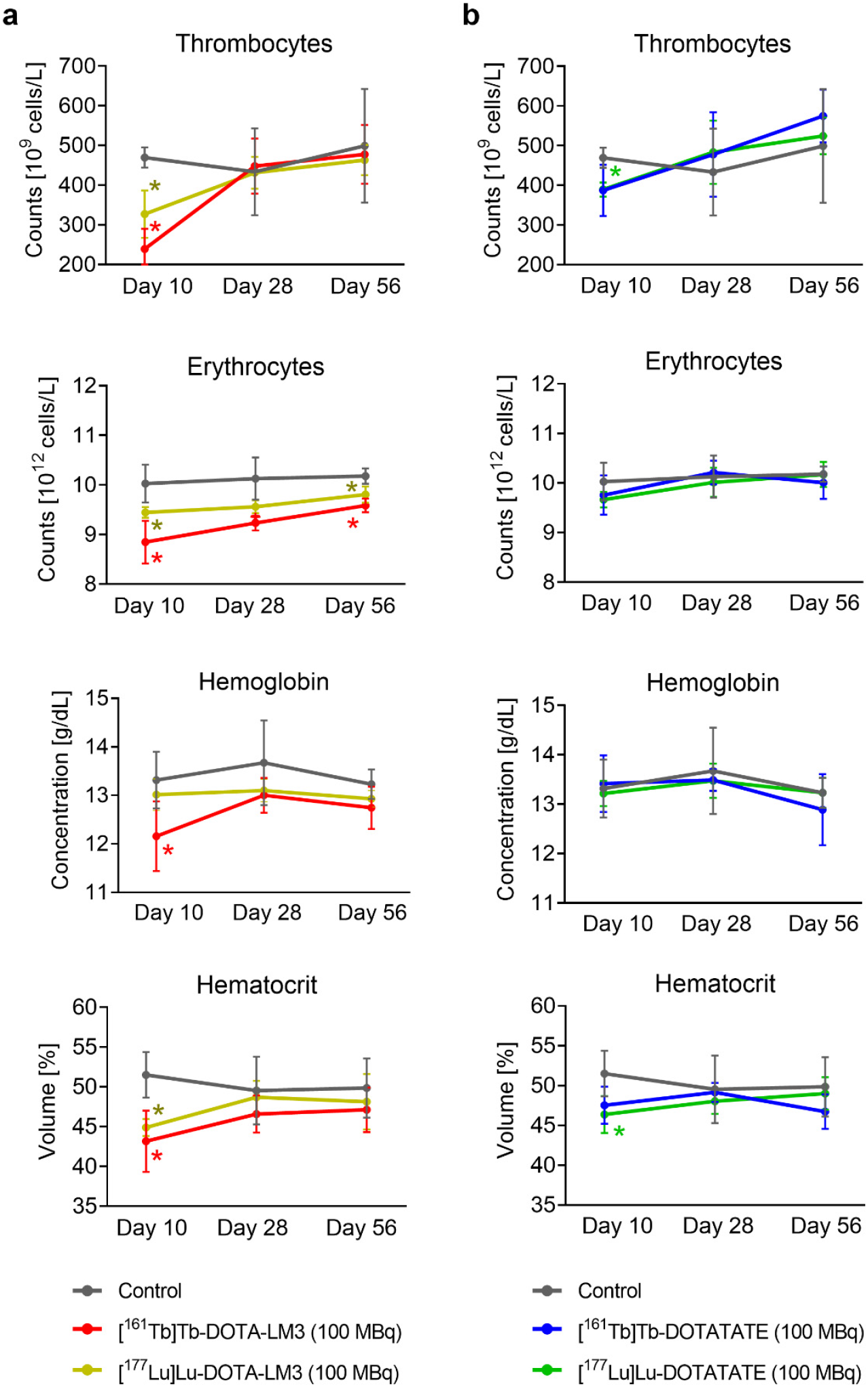
**a/b** Thrombocyte and erythrocyte counts as well as hemoglobin and hematocrit values on Day 10, Day 28 and Day 56 after treatment of mice with 100 MBq (1 nmol); **a** [^161^Tb]Tb-/[^177^Lu]Lu-DOTA-LM3; **b** [^161^Tb]Tb-/[^177^Lu]Lu-DOTATATE in comparison to untreated control mice. (*) Values that differed significantly (*p*<0.05) from those of the control group.

On Day 28, a decrease of 35−55% was determined for leukocyte counts in treated mice compared to untreated controls (*p*<0.05; Fig. 5; Supplementary Material, Table S11). At study end, mice treated with ^177^Lu-based radiopeptides had average leukocyte counts comparable to those of the control group ((8.7 ± 1.8) × 10^9^ cells/L; *p*>0.05). This was not the case for mice treated with [^161^Tb]Tb-DOTA-LM3 or [^161^Tb]Tb-DOTATATE for which leukocytes were still about 30% lower ((6.0 − 6.2) × 10^9^ cells/L; *p*>0.05) than control values at study end. These changes were ascribed to the drop in lymphocytes, whereas the neutrophils remained in the same range for treated and untreated control mice (Fig. 5).

**Fig. 5.**
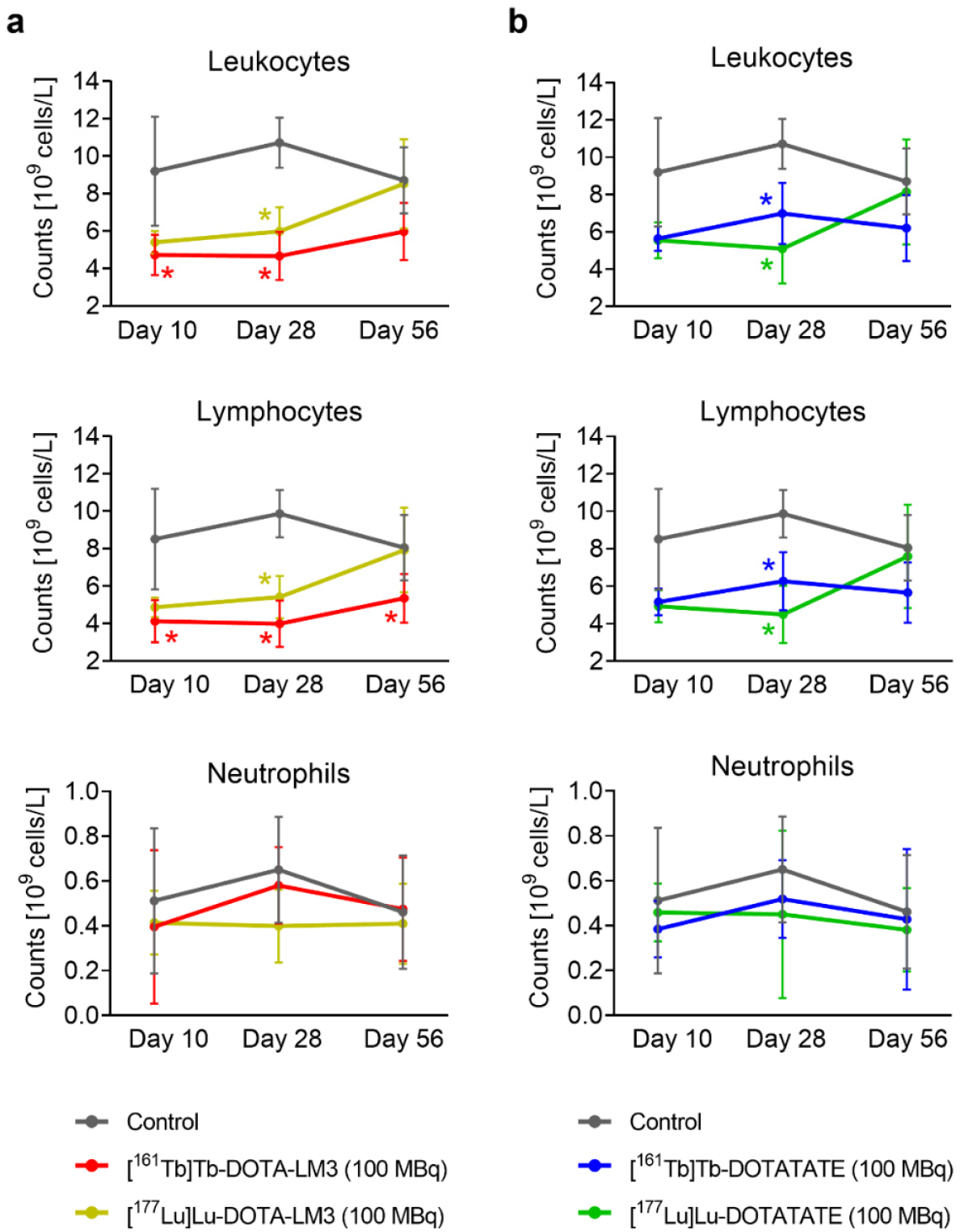
**a/b** Leukocyte, lymphocyte and neutrophil counts on Days 10, 28 and 56 after treatment of mice with 100 MBq (1 nmol); **a** [^161^Tb]Tb-/[^177^Lu]Lu-DOTA-LM3; **b** [^161^Tb]Tb-/[^177^Lu]Lu-DOTATATE in comparison to untreated control mice. (*) Values that differed significantly (*p*<0.05) from those of the control group.

No morphologically abnormal cells were present in the blood smears of any of the investigated mice, however, on Day 10, fewer nucleated cells were observed in all treated mice (Supplementary Material, Tables S12). From Day 28 on, many of these animals showed a premature release of precursor and immature cells from the myeloid lineage in the peripheral blood. Fewer reticulocytes were visible on blood smears of treated mice than on those of untreated control mice.

Bone marrow smears of treated mice demonstrated increased myelopoiesis and decreased erythropoiesis compared to samples obtained from control mice (Supplementary Material, Table S13). In treated mice, an increased number of hematopoietic precursor cells was observed, wherein many of these were of neutrophilic lineage. On the other hand, the decrease in erythropoiesis was attributed to the reduced number of premature erythrocyte precursors cells (Supplementary Material).

The average blood urea nitrogen levels of treated mice (7.8−8.9 mM) were slightly higher than those of control mice (6.3 ± 3.2 mM; *p*>0.05) while alkaline phosphatase levels were only marginally lower for treated mice (88-108 U/L) than for controls (115 ± 23 U/L; *p*>0.05). Albumin levels were comparable for all groups of mice (20-22 g/L, *p*>0.05; Supplementary Material, Table S14).

Frequent, but mild treatment-related abnormalities were observed in tissue sections of the spleen, kidneys and liver. The severity of lesions was similar among the treatment groups (Supplementary Material, Fig. S6 and S7, Table S15). Minimal lymphoid hyperplasia was present in the spleen of all treated mice, while this lesion was not seen in any of the control mice (Supplementary Material, Fig. S6). Subcapsular dilatation of blood vessels was observed in 6 out of 7 mice that received [^161^Tb]Tb-DOTA-LM3 and in individual mice of the other treatment groups, but was completely absent in control mice (Fig. 6a/b; Supplementary Material, Fig. S7).

**Fig. 6.**
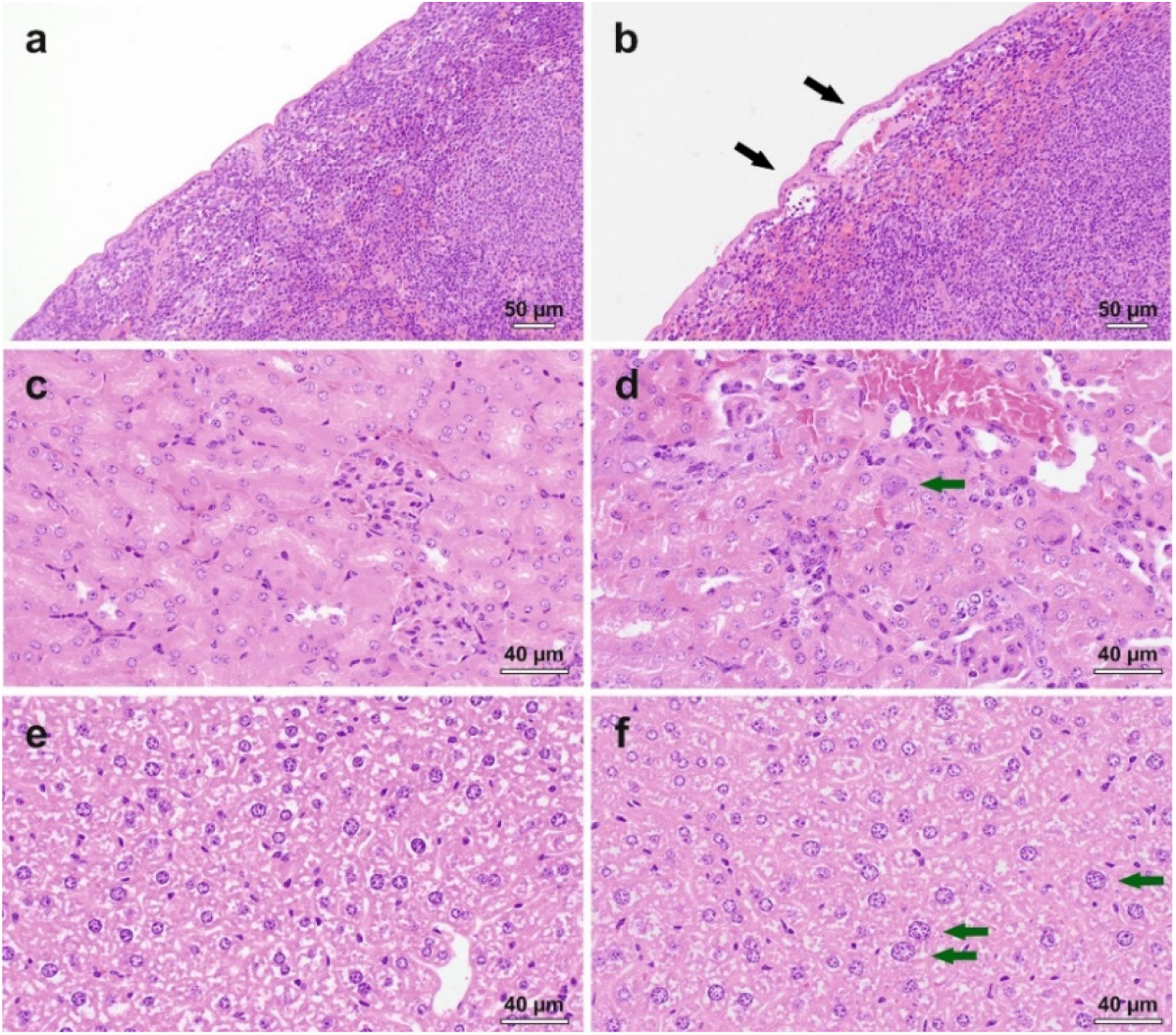
**a-f** Representative images showing treatment-related histopathological findings. **a** Spleen of an untreated control mouse; **b** spleen of a treated mouse with subcapsular dilatation of the blood vessel. **c** kidney of an untreated control mouse; **d** kidney of a treated mouse with karyomegaly. **e** liver of an untreated control mouse; **f** liver of a treated mouse with karyomegaly. Black arrows: subcapsular dilatation; green arrows: karyomegaly.

Enlarged nuclei (karyomegaly) in epithelial cells of the proximal tubule were present in all mice injected with [^161^Tb]Tb-DOTATATE or [^177^Lu]Lu-DOTATATE, while this was not seen in kidneys of control mice (Fig. 6c/d; Supplementary Material, Fig. S7). This was also observed in one mouse injected with [^161^Tb]Tb-DOTA-LM3 and five mice that received [^177^Lu]Lu-DOTA-LM3. Karyomegaly was also observed in hepatocytes of all groups of treated mice, but with the least incidence in the group treated with [^161^Tb]Tb-DOTA-LM3 (Fig. 6e/f; Supplementary Material, Fig. S7).

## Discussion

The postulated increased absorbed dose to single cancer cells when using terbium-161 instead of lutetium-177 during PRRT [15, 17] may lead to a more effective treatment outcome, as previously demonstrated in preclinical studies [19]. The question arises, however, whether specific binding of somatostatin analogues to the few SSTR-positive cells in the bone marrow (CD34^+^ hematopoietic stem cells and early lineage progenitor cells [25]) would enhance adverse events of ^161^Tb-based PRRT compared to the application of [^177^Lu]Lu-DOTATATE [30].

The in vitro data obtained from isolated murine bone marrow cells confirmed minute, but specific binding of somatostatin analogues and, hence, the presence of a small fraction of SSTR-expressing cells. This was corroborated by the data of simulation studies, in which similar patterns of relative binding of the radiopeptides were obtained in a mixture of tumor cells comprising ≤10% of SSTR-positive cells. Moreover, higher accumulation of the radiopeptides was found in the femur than in the blood, which was not commonly the case for tumor-targeting agents such as [^177^Lu]Lu-PSMA-617 [31], indicating again the specific accumulation of the somatostatin analogues in bone marrow cells as previously proposed by others [32, 33].

Despite the calculated 3.2-fold increased absorbed bone marrow dose for the SSTR antagonist than for the agonist, blood cell counts were equally affected with only mild and transient changes in lymphocyte and thrombocyte counts for either peptide type applied at 20 MBq/mouse (Study I). This obviously safe activity level, which would translate to about 4.5 GBq per patient [21], was previously found to be effective in delaying tumor growth in mice [10, 19, 34].

Effects following the injection of the radiopeptides at 100 MBq per mouse were investigated to determine differences between the two somatostatin analogues and the two radionuclides, respectively. Under these conditions, more pronounced, but mostly transient, changes of blood cell counts were determined. Fewer lymphocytes were present in peripheral blood after treatment with [^161^Tb]Tb-DOTA-LM3 and [^161^Tb]Tb-DOTATATE than with the ^177^Lu-labeled counterparts with a nadir on Day 28. Recovery was seen by study end, even though the quantitative analysis of the bone marrow smears revealed only few mature lymphocytes. The fast recovery may, thus, be ascribed to the compensatory lymphoid hyperplasia in the spleen, which is unique in rodents [35]. This could explain the increased resistance of mice towards high bone marrow doses as compared to human patients. The unaltered number of prolymphocytes in the bone marrow was, however, an indication that lymphocyte counts would recover over time irrespective of this splenic mechanism.

Effects on erythrocytes and thrombocytes appeared to be more pronounced after the treatment with the SSTR antagonist than with the agonist, irrespective of the employed radionuclide. This observation was in line with data of early phase clinical trials with ^177^Lu-labeled SSTR antagonists that showed more frequent thrombocytopenia than commonly seen after application of SSTR agonists, while lymphocytopenia was similar in patients treated with SSTR antagonists and SSTR agonists [6, 13, 14]. In mice, a treatment-related effect on erythropoiesis was also reflected by less reticulocytes in the peripheral blood, however, reduced erythropoiesis appeared to be of minimal concern since the well-being of the mice was not seriously affected. Although the rapid recovery of erythrocyte could be due to the hematopoietic function of the spleen in mice, the presence of blast cells in the bone marrow suggests that erythrocytes would have recovered over time anyway.

Effects related to SSTR-expressing bone marrow cells could eventually be reduced by adjustment of the injected peptide mass to saturate the receptors as previously demonstrated [36, 37]. In Study II, a relatively high peptide amount of 1 nmol per mouse was applied, which may have partially saturated SSTRs in bone marrow cells, as previously shown by Borgna et al. [19]. In the clinical setting, it was also demonstrated that the applied peptide mass critically affects the uptake of somatostatin analogues in normal organs [38, 39] and, hence, this could be a relevant aspect for the design of future clinical applications [12].

Histological investigations revealed subcapsular dilatation of blood vessels in the spleen, which occurred in a dose-dependent manner and was most frequently observed in mice treated with [^161^Tb]Tb-DOTA-LM3. Potential consequences of this impairment remained unexplored in this study. In Study II, the absorbed kidney dose in mice injected with [^161^Tb]Tb-DOTATATE (30 Gy) or [^177^Lu]Lu-DOTATATE (22 Gy) were within a reasonable range, which was, however, not the case for [^161^Tb]Tb-DOTA-LM3 (100 Gy) and [^177^Lu]Lu-DOTA-LM3 (76 Gy). Previously, a renal dose of 23-30 Gy was found to be safe, however, above this limit, damage to the kidneys was observed 3-4 months after treatment with folate radioconjugates [40, 41]. Tubular basophilia and karyomegaly of minimal severity in the kidneys and slightly increased blood urea nitrogen levels in blood plasma were indeed seen in treated mice, however, these events were not more frequently observed after application of the SSTR antagonist than after injection of the agonist. The liver mass of treated mice was higher in comparison to control values, but the alkaline phosphatase levels in plasma remained in the normal range. Histological investigations revealed karyomegaly in the liver of treated animals, although the frequency and severity of this lesion did not correlate with the absorbed radiation dose. Overall, the findings referring to kidneys and liver were not of concern, despite the extremely high activity that was applied. Conclusions about long-term impairment of renal and hepatic functions should be based on studies over at least six months [40].

Limitations of this study refer to the fact that blood plasma parameters, indicative for kidney and liver function, were not determined earlier than Day 56 after radiopeptide application. Moreover, the study was performed with healthy mice and, hence, not reflecting the patient situation accurately. The focus was more on the effects on normal tissue related to radioligand therapy, while excluding potential impairment of the well-being of mice due to the tumor burden. It has to be critically noted that the transferability to the situation in humans remains unclear, although analogies based on the data obtained with the clinically-established [^177^Lu]Lu-DOTATATE as well as with [^177^Lu]Lu-DOTA-LM3 will certainly help clinicians to judge on the safety of the ^161^Tb-based analogues.

## Conclusion

In this preclinical study, no major hematological changes were observed after PRRT in mice at an activity which would translate to the currently used activity per treatment cycle in clinics. In mice treated with radiopeptides at a five-fold higher activity, the blood cell counts were more affected for DOTA-LM3 than for DOTATATE while the replacement of lutetium-177 with terbium-161 had only minor implications. The drop in blood cell counts was mostly transient and histological abnormalities of minimal severity, hence, these effects will most probably prove clinically insignificant. Based on this study, the application of [^161^Tb]Tb-DOTA-LM3 and [^161^Tb]Tb-DOTATATE should be safe at activity levels that are currently recommended for the ^177^Lu-based analogues.

## Acknowledgment

The authors thank Damian Wild and Korbinian Krieger for proof-reading of the manuscript and helpful advice; they thank the team at ILL for the production of terbium-161, Pascal V. Grundler and Colin Hillhouse for the separation of terbium-161 at PSI and Susan Cohrs and Fan Sozzi-Guo for technical assistance during the in vitro and in vivo experiments at PSI.

## Funding Information

Sarah D. Busslinger and Ana Katrina Mapanao were funded by a SNSF grant (N° 320030_188978; PI: Cristina Müller); Ana Katrina Mapanao received funding from the European Union’s Horizon 2020 research and innovation program under the Marie Skłodowska-Curie grant agreement N^o^ 884104. The research was further supported by the Ulrich Peter & Hans Rudolf Wirz Foundation via Swiss Cancer Research (KFS-5624-08-2022; PI: Cristina Müller). Peter Bernhardt received grants from the Swedish Cancer Society, the King Gustav V Jubilee Clinic Cancer Research Foundation, the Swedish state under the agreement between the Swedish government and the county councils, the ALF-agreement. ITM Isotope Technologies Munich GmbH, Germany, provided ^177^Lu for the reported studies and funded the histological analyses performed by AnaPath Services GmbH.

## Ethics approval

This study was performed in agreement with the national law and PSI-internal guidelines of radiation safety protection. In vivo experiments were approved by the local veterinarian department and ethics committee and conducted in accordance with the Swiss law of animal protection.

## Conflict of interests

The University Hospital Basel and the Paul Scherrer Institute have filed a patent with regard to [^161^Tb]Tb-DOTA-LM3. R. Schibli, N. van der Meulen and C. Müller are listed as co-inventors on the patent application.

## Notes

### Competing Interest Statement

The University Hospital Basel and the Paul Scherrer Institute have filed a patent with regard to [161Tb]Tb-DOTA-LM3. R. Schibli, N. van der Meulen and C. Mueller are listed as co-inventors on the patent application.

### Summary of Updates

Title adapted; Discussion section updated and limitations of the study included; Supplemental file updated with additional data on autoradiography studies.

