## Supplementary Material for "Comparison of the tolerability of ^161^Tb- and ^177^Lu-labeled somatostatin analogues in the preclinical setting"

### 1. Terbium-161 and lutetium-177

***Purpose:*** Terbium-161 and lutetium-177 were used for labeling of the somatostatin analogues to enable the dosimetry and tolerability studies in the preclinical setting.

***Methods:*** Terbium-161 was obtained from the Radionuclide Development Group at the Paul Scherrer Institute. It was produced by neutron irradiation of enriched gadolinium-160 targets (Isoflex, San Francisco, United States) at the SAFARI-1 reactor at Necsa, Pelindaba, South Africa, or at the RHF reactor at the Institut Laue Langevin, Grenoble, France using the $^{160}Gd(n,\gamma)^{161}Gd \rightarrow ^{161}Tb$ nuclear reaction. After chemical separation performed as previously described [1], no-carrier-added (n.c.a.) [$^{161}$Tb]$TbCl_3$ was obtained in 0.05 M HCl. In addition, n.c.a. [$^{161}$Tb]$TbCl_3$ was purchased from Terthera (Breda, the Netherlands) via Solumedics AG (Aarau, Switzerland). Lutetium-177 was obtained as n.c.a. [$^{177}$Lu]$LuCl_3$ in 0.04 M HCl from ITM Medical Isotopes GmbH, Germany. Terbium-161 was measured using a dose calibrator (NUVIA Instruments GmbH, ISOMED 2010) which was calibrated for variable vial geometries and filling volumes by the Institute of Radiation Physics, Lausanne, Switzerland [2]. The linear range for the measurement of terbium-161 in the γ-counter (PerkinElmer, Wallac Wizard 1480, MA, United States) was assessed using dilution series. The same procedure was performed for lutetium-177.

***Results:*** The calibrated ionization chamber allowed accurate measurement of terbium-161 samples in various vials and volumes. Accurate measurement of lutetium-177 was enabled through the standard settings of this same dose calibrator. The linear range of the γ-counter to measure terbium-161 was defined from 100 counts per minute (cpm) to $2 \times 10^6$ cpm. The linear range for the measurement of lutetium-177 was defined from 200 cpm to $8 \times 10^6$cpm. The impact of the sample volume on the counts detected by the γ-counter was found to be negligible for the range of relevant volumes used in this study.

### 2. Radiolabeling of the somatostatin analogues

***Purpose:*** DOTA-LM3 and DOTATATE were radiolabeled with terbium-161 or lutetium-177 for the preclinical studies.

***Methods***: A stock solution of DOTA-LM3 was prepared using MilliQ water to obtain a final peptide concentration of 1 mM. The stock solution of DOTATATE at the same concentration was prepared using a mixture of MilliQ water and 25% dimethylsulfoxide. Radiolabeling of DOTA-LM3 and DOTATATE with terbium-161 or lutetium-177 was carried out under standard labeling conditions at pH ~4.5 using a 1:5 (*v*/*v*) mixture of sodium acetate (0.5 M) and HCl (0.05 M) as previously reported [3]. The reaction mixtures were incubated for 10 min at 95 °C. Quality control of the radiolabeled peptides was performed by reversed-phase high performance liquid chromatography (HPLC) using an aliquot of the reaction mixture diluted in MilliQ water containing penta-sodium diethylenetriamine pentaacetic acid ($Na_5$-DTPA; 50 μM). The HPLC system (Merck Hitachi LaChrom HPLC) was equipped with a D-7000 interface, a L-7200 autosampler, a radioactivity detector (LB 506 B; Berthold) and a L-7100 pump connected to a C18 column (Xterra$^{TM}$, 5 μm, 4.6×150 mm, Waters, Milford, MA, U.S.A). A linear gradient of acetonitrile (5–

80%) and MilliQ water with 0.1% trifluoroacetic acid (95–20%) was applied over 15 min as a mobile phase using a flow rate of 1 mL/min.

***Results:*** The radiopeptides were obtained with a radiochemical purity of >98% at a molar activity of up to 100 MBq/nmol (Fig. S1). The radiopeptides were, thus, employed without further purification steps for the herein reported preclinical investigations.

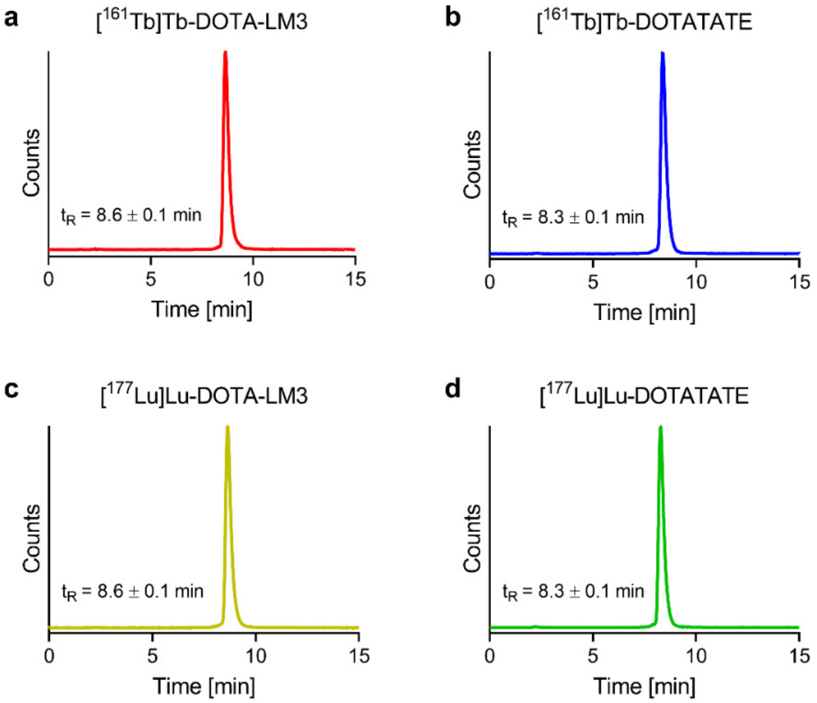


**Fig. S1 a–d** Representative HPLC chromatograms of the herein applied radiopeptides prepared at 100 MBq/nmol. **a** Chromatogram of [$^{161}$Tb]Tb-DOTA-LM3; **b** Chromatogram of [$^{161}$Tb]Tb-DOTATATE; **c** Chromatogram of [$^{177}$Lu]Lu-DOTA-LM3 and **d** Chromatogram of [$^{177}$Lu]Lu-DOTATATE (Traces of unreacted radionuclide appeared with a retention time ($t_R$) of 2.2 min.).

### 3. Biodistribution studies

***Purpose:*** The tissue distribution of [$^{161}$Tb]Tb-DOTA-LM3 and [$^{161}$Tb]Tb-DOTATATE was assessed in immunocompetent FVB mice to enable dosimetry estimations for normal tissues and organs.

***Methods:*** FVB mice were acclimatized for at least one week before initiation of the in vivo studies. After intravenous injection of [$^{161}$Tb]Tb-DOTA-LM3 (5 MBq, 1 nmol) or [$^{161}$Tb]Tb-DOTATATE (5 MBq, 1 nmol) diluted in 100 μL phosphate-buffered saline (PBS pH 7.4) containing 0.05% bovine serum albumin (BSA), the mice were dissected 2 h, 6 h, 24 h or 72 h post injection (p.i.). Selected organs and tissues were collected, weighed and counted for activity using a γ-counter (PerkinElmer, Wallac Wizard 1480, Waltham, MA, United States). Defined volumes of the injection solution were measured at the same time to obtain decay-corrected data. The injected activity per gram tissue mass (% IA/g) was calculated as the average ± standard deviation (SD) of n=3 mice per timepoint.

***Results:*** The results are reported in the main article and listed in Tables S1 and S2.

**Table S1** Biodistribution data obtained in immunocompetent FVB mice at various time points after injection of [$^{161}$Tb]Tb-DOTA-LM3. Decay-corrected data of accumulated activity are shown as % IA/g tissue, representing the average ± SD.

| | [$^{161}$Tb]Tb-DOTA-LM3 | | | |
|---|---|---|---|---|
| **Organ** | **2 h p.i.** | **6 h p.i.** | **24 h p.i.** | **72 h p.i.** |
| | n=3 | n=3 | n=3 | n=3 |
| | [% IA/g] | [% IA/g] | [% IA/g] | [% IA/g] |
| Blood | 0.13 ± 0.02 | 0.03 ± 0.01 | 0.01 ± 0.01 | <0.01 |
| Heart | 0.12 ± 0.02 | 0.09 ± 0.01 | 0.05 ± 0.01 | 0.02 ± 0.01 |
| Femur | 0.43 ± 0.04 | 0.33 ± 0.04 | 0.26 ± 0.02 | 0.16 ± 0.02 |
| Spleen | 0.26 ± 0.01 | 0.21 ± 0.01 | 0.13 ± 0.01 | 0.08 ± 0.01 |
| Thymus | 0.65 ± 0.11 | 0.55 ± 0.03 | 0.43 ± 0.07 | 0.29 ± 0.02 |
| Kidneys | 17 ± 2 | 19 ± 1 | 8.3 ± 0.6 | 3.2 ± 0.2 |
| Adrenals | 0.55 ± 0.07 | 0.52 ± 0.02 | 0.35 ± 0.15 | 0.16 ± 0.02 |
| Pancreas | 3.8 ± 0.1 | 3.1 ± 0.2 | 1.6 ± 0.2 | 0.42 ± 0.03 |
| Stomach | 3.2 ± 0.3 | 2.6 ± 0.4 | 1.8 ± 0.2 | 0.86 ± 0.08 |
| Lung | 2.3 ± 0.3 | 1.7 ± 0.2 | 1.0 ± 0.04 | 0.50 ± 0.04 |
| Liver | 0.66 ± 0.07 | 0.62 ± 0.01 | 0.36 ± 0.01 | 0.17 ± 0.01 |
| Intestines | 0.30 ± 0.01 | 0.25 ± 0.02 | 0.13 ± 0.01 | 0.06 ± 0.01 |
| Muscle | 0.06 ± 0.01 | 0.04 ± 0.01 | 0.02 ± 0.01 | 0.01 ± 0.01 |
| Brain | 0.02 ± 0.01 | 0.03 ± 0.01 | 0.02 ± 0.01 | 0.01 ± 0.01 |
| Salivary glands | 0.15 ± 0.01 | 0.12 ± 0.01 | 0.06 ± 0.01 | 0.03 ± 0.01 |

**Table S2** Biodistribution data obtained in immunocompetent FVB mice at various time points after injection of [$^{161}$Tb]Tb-DOTATATE. Decay-corrected data of accumulated activity are shown as % IA/g tissue, representing the average ± SD.

| | [$^{161}$Tb]Tb-DOTATATE | | | |
|---|---|---|---|---|
| **Organ** | **2 h p.i.** | **6 h p.i.** | **24 h p.i.** | **72 h p.i.** |
| | n = 3 | n = 3 | n = 3 | n = 3 |
| | [% IA/g] | [% IA/g] | [% IA/g] | [% IA/g] |
| Blood | 0.04 ± 0.01 | 0.01 ± 0.01 | 0.01 ± 0.01 | <0.01 |
| Heart | 0.04 ± 0.01 | 0.03 ± 0.01 | 0.01 ± 0.01 | 0.01 ± 0.01 |
| Femur | 0.20 ± 0.01 | 0.14 ± 0.02 | 0.10 ± 0.01 | 0.05 ± 0.01 |
| Spleen | 0.09 ± 0.01 | 0.06 ± 0.01 | 0.04 ± 0.01 | 0.02 ± 0.01 |
| Thymus | 0.69 ± 0.08 | 0.59 ± 0.06 | 0.35 ± 0.03 | 0.20 ± 0.02 |
| Kidneys | 9.0 ± 0.6 | 6.7 ± 1.3 | 3.4 ± 0.5 | 1.2 ± 0.4 |
| Adrenals | 0.52 ± 0.06 | 0.36 ± 0.14 | 0.30 ± 0.06 | 0.18 ± 0.04 |
| Pancreas | 2.4 ± 0.1 | 1.1 ± 0.2 | 0.28 ± 0.03 | 0.07 ± 0.01 |
| Stomach | 2.1 ± 0.4 | 1.4 ± 0.1 | 0.79 ± 0.18 | 0.38 ± 0.04 |
| Lung | 1.4 ± 0.2 | 0.91 ± 0.17 | 0.51 ± 0.08 | 0.25 ± 0.03 |
| Liver | 013 ± 0.01 | 0.09 ± 0.01 | 0.05 ± 0.01 | 0.03 ± 0.01 |
| Intestines | 0.25 ± 0.02 | 0.14 ± 0.01 | 0.08 ± 0.02 | 0.04 ± 0.01 |
| Muscle | 0.02 ± 0.01 | 0.01 ± 0.01 | 0.01 ± 0.01 | <0.01 |
| Brain | 0.01 ± 0.01 | 0.02 ± 0.01 | 0.01 ± 0.01 | 0.01 ± 0.01 |
| Salivary glands | 0.07 ± 0.01 | 0.04 ± 0.01 | 0.02 ± 0.01 | 0.01 ± 0.01 |

### 4. In vitro autoradiography using bone marrow cells

***Purpose:*** The binding of [$^{177}$Lu]Lu-DOTA-LM3 and [$^{177}$Lu]Lu-DOTATATE to isolated murine bone marrow cells was assessed and compared in vitro.

***Methods:*** Bone marrow cells were isolated according to a previously published protocol [4]. The femurs, tibiae and iliac bones from immunocompetent FVB mice were collected and muscle tissue was carefully removed. The bones were put into ice-cold PBS until further processed. The bones were cut open at each end using a scalpel and placed in 200-µL Eppendorf pipette tips of which the end was cut to enlarge the opening. The pipette tips with the bones were placed in Eppendorf tubes (filled with 100 µL RPMI cell culture medium) followed by centrifugation (15000 rcf) for 20 s to isolate the bone marrow cells. The centrifugation step was repeated using another Eppendorf tube (filled with 100 µL RPMI cell culture medium), to which the pipette tip with the bones was transferred. Afterwards, the collected bone marrow cells were resuspended and pooled. Filtration of the cells using a cell strainer (70 µm pore size) was followed by counting the bone marrow cells manually to prepare cell suspensions diluted in RPMI cell culture medium. Samples of 5 × 10$^6$ bone marrow cells in 400 µL RPMI culture medium were prepared in Eppendorf tubes to perform binding experiments with both radiopeptides. Cell culture medium without unlabeled peptide (100 µL) or medium containing various amounts of unlabeled peptide (0.5, 2.5 or 5 pmol in 100 µL RPMI medium) were added and the samples were incubated for 10 min at 37 °C while shaking. The respective radiopeptides were prepared at 100 MBq/nmol and diluted in RPMI medium at a concentration of 0.1 MBq/mL. A volume of 500 µL of this radiopeptide solution was added to the bone marrow samples to obtain final volumes of 1 mL. This means that the samples contained 0.5 pmol radiopeptide of an activity of 50 kBq only or additionally a 1-fold, 5-fold or 10-fold excess of unlabeled peptide relative to the $^{177}$Lu-labeled radiopeptide. The samples were incubated for 2 h at 37 °C while shaking. Afterwards, the samples were centrifuged (10000 rcf) for 5 min at 4 °C. The cell pellet was washed 3 times with 1 mL ice-cold PBS followed by resuspension in 25 µL PBS. The bone marrow cell samples were put on a membrane and let to dry (Fig. S2).

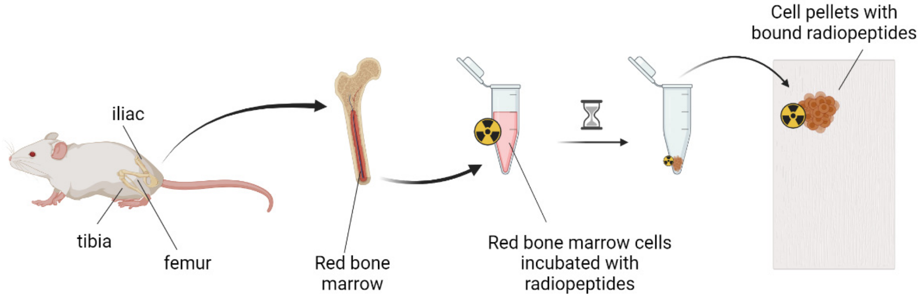


**Fig. S2** Schematic outline of the procedure to evaluate the binding of radiopeptides to isolated bone marrow cells (image created with BioRender.com).

The membrane with the bone marrow cell samples was exposed to a super resolution phosphor imaging screen (SR 7001486, PerkinElmer, Waltham, MA, United States) for 72 h. Images were obtained using a storage phosphor system (Cyclone Plus, PerkinElmer™, Waltham, MA, United States) and the signals were quantified using OptiQuant software (version 5.0, Bright Instrument Co Ltd., PerkinElmer, Waltham, MA, United States). The data were presented as average ± SD of 3 independent experiments, each performed in duplicates.

The experimental setting was validated with regard to the pattern that can be expected when using a cell mixture with a small fraction of SSTR-positive cells as it is reported to be the case in the bone marrow [5]. For this purpose, the same assay was performed with SSTR-negative KB tumor cells mixed with various fractions of SSTR-positive AR42J tumor cells (10%, 1% or 0.1%). Quantification of the signal intensities were performed after a 10-minute exposure to the phosphor screen. The data were presented as average ± SD of at least 2 independent experiments, each performed in duplicates.

***Results:*** The results of the binding of the radiopeptides to isolated bone marrow cells are reported in the main article. Data based on the simulation study performed with mixtures of KB and AR42J tumor cells revealed a signal intensity that correlated positively with the fraction of SSTR-positive AR42J cells in the tumor cell mixture (Fig. S3a). Moreover, the signal intensity was higher for the SSTR antagonist than for the SSTR agonist. The pattern of SSTR-blocking with the same amount of excess unlabeled peptide was similar as that obtained with bone marrow cells (Fig. S3b/c), which clearly confirmed the presence of SSTR-positive cells present in the bone marrow. On the other hand, the intensity of the signal was still much higher for the tumor cell mixtures than for bone marrow cells which were exposed to the phosphor screen for 10 min versus 72 h, respectively.

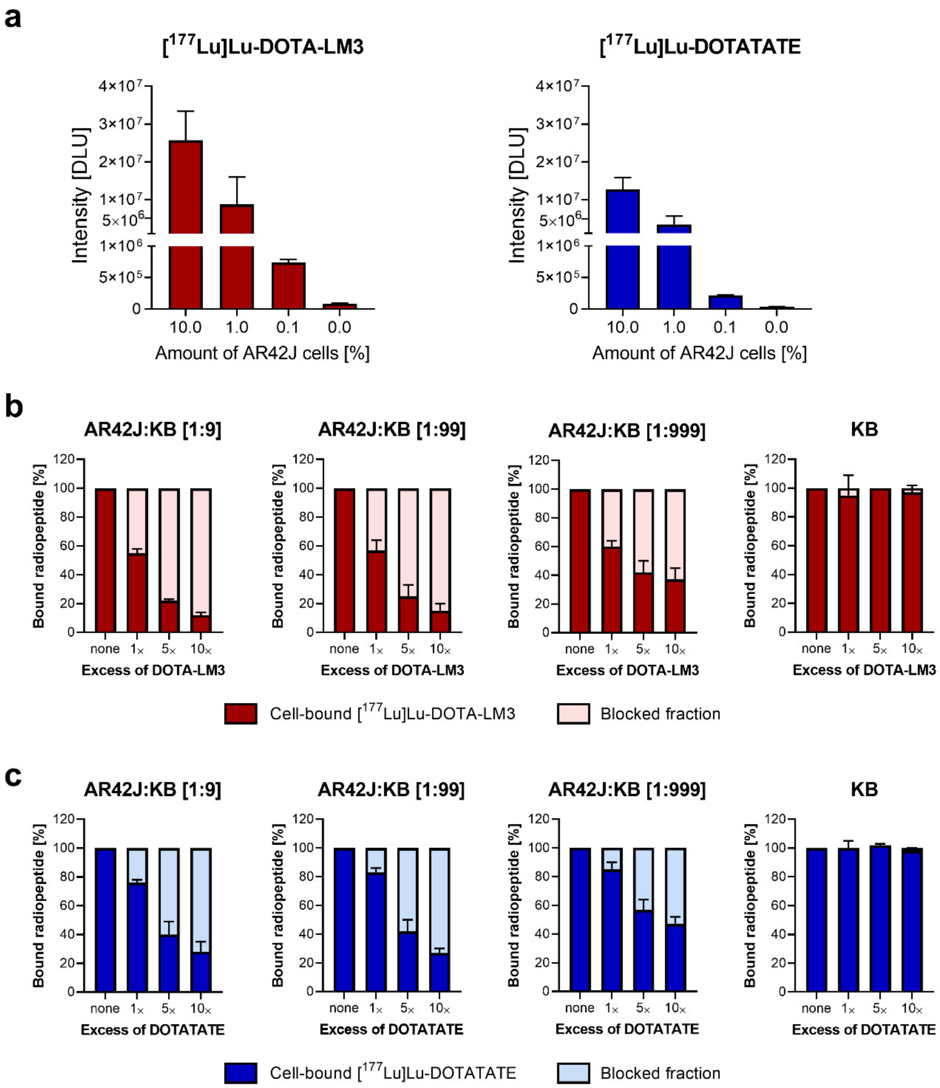


**Fig. S3** **a** Comparison of the signal intensities indicated as DLU for various mixtures of SSTR-negative KB tumor cells and SSTR-positive AR42J tumor cells. **b/c** Binding of the radiopeptides ([$^{177}$Lu]Lu-DOTA-LM3 (**b**) and [$^{177}$Lu]Lu-DOTATATE (**c**)) to various mixtures of AR42J and KB cells incubated in the absence and presence of excess unlabeled peptide. The binding of the radiopeptides to these cell mixtures is reported relative to the total binding of the radiopeptides without unlabeled peptide (set as 100%).

### 5. Dosimetry calculations

***Purpose:*** The absorbed organ doses were estimated based on the non-decay-corrected biodistribution data obtained with immunocompetent FVB mice after injection of [$^{161}$Tb]Tb-DOTA-LM3 or [$^{161}$Tb]Tb-DOTATATE. Data for [$^{177}$Lu]Lu-DOTA-LM3 and [$^{177}$Lu]Lu-DOTATATE were calculated based on the assumption that the tissue distribution profiles remained the same, irrespective of the employed radiolanthanide.

***Methods:*** The absorbed dose in various organs and tissues was based on tissue distribution data obtained from mouse studies performed with [$^{161}$Tb]Tb-DOTA-LM3 and [$^{161}$Tb]Tb-DOTATATE, which were re-calculated to obtain non-decay-corrected biodistribution data (Tables S1 and S2). The activity concentration was plotted against the time and bi-exponential curve fits were applied. The calculations were performed for all possible combinations of the data points, resulting in a total of 81 combinations with n=3 mice for each of the 4 time points. Each curve fit was integrated to infinity to obtain the time integrated activity concentration (TIAC). The absorbed energy per decay was obtained from the mouse phantom MOBY [6]. Although the enhanced activity uptake in the femur indicated a specific uptake in the bone marrow as described by Hemmingsson et al. [7], it was not feasible to experimentally isolate the bone marrow and measure the activity concentration therein. Instead, the mass relation between the cortical bone and bone marrow in the femur was used to estimate the activity concentration in the bone marrow, assuming that the activity concentration in the cortical bone and muscle was equal. The masses were obtained from the MOBY phantom (19.8 mg and 16.9 mg for left and right cortical bone of femur, and 20.1 mg and 19.3 mg for bone marrow content in left and right femur, respectively). The absorbed doses were calculated by multiplying the TIAC with the corresponding absorbed energy for all tissues. It was assumed that the tissue distribution of [$^{161}$Tb]Tb-DOTA-LM3 and [$^{161}$Tb]Tb-DOTATATE is equal to that of the respective $^{177}$Lu-labeled counterparts, hence, the absorbed doses for [$^{177}$Lu]Lu-DOTA-LM3 and [$^{177}$Lu]Lu-DOTATATE was calculated by multiplying the TIAC (adjusted for the difference in physical half-life between $^{161}$Tb and $^{177}$Lu) and the absorbed energy for all tissues.

***Results:*** The results are presented and discussed in the main article.

### 6. Study I: Tolerability of [$^{161}$Tb]Tb-DOTA-LM3 and [$^{161}$Tb]Tb-DOTATATE (20 MBq/mouse)

The tolerability of 20 MBq [$^{161}$Tb]Tb-DOTA-LM3 was assessed in mice and compared to that of 20 MBq [$^{161}$Tb]Tb-DOTATATE. This activity was chosen in Study I because it would translate in ~4.5 GBq for human patients based on the ratio of the body mass between a human patient and a mouse (70 kg/0.025 kg) and their surfaces (1/12.3) [8].

#### 6.1. Body masses, organ masses and respective organ-to-brain mass ratios

***Purpose:*** The body mass of mice was monitored during the entire study over 56 days as it presents a relevant parameter for the general health status of the mice. At study end, potential changes in the organ masses and organ-to-brain mass ratios of mice treated with [$^{161}$Tb]Tb-DOTA-LM3 or [$^{161}$Tb]Tb-DOTATATE were assessed in comparison to untreated control mice.

***Methods:*** The body mass of mice was measured three times a week and expressed as the relative body mass to the initial body mass at Day 0 ($BM_0$), defined as [$BM_x$/$BM_0$], where $BM_x$ is the body mass in grams on a given Day x. Endpoint criteria were determined based on loss of body mass and signs of unease and pain using a predefined scoring system considering several parameters including behavior and appearance, among others. Immediately after euthanasia at study end on Day 56, the brain, spleen, kidneys and liver were collected and weighed to calculate the respective organ-to-brain mass ratios.

***Results:*** The results are reported mainly in the main article. The body mass of mice increased continuously over the course of the study in all groups of mice (Fig. S4).

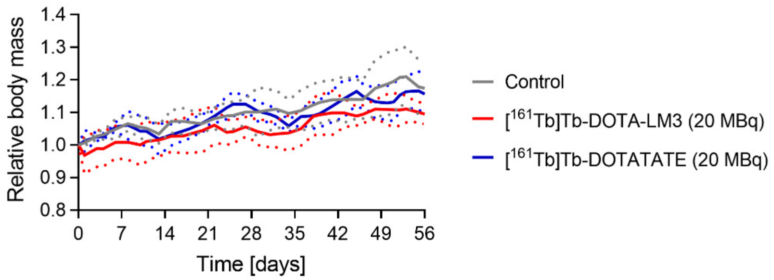


**Fig. S4** Body mass of mice treated with 20 MBq [$^{161}$Tb]Tb-DOTA-LM3 or 20 MBq [$^{161}$Tb]Tb-DOTATATE in comparison to the body mass of untreated control mice. The body mass was monitored over the entire period of the study which lasted for two months.

Organ masses of treated mice did not significantly differ from those of untreated controls, however, a trend of increased liver masses in treated mice was observable (Table S3). The liver-to-brain mass ratios were not affected in the case of mice treated with [$^{161}$Tb]Tb-DOTA-LM3, however, a significantly increased ratio was found for mice treated with [$^{161}$Tb]Tb-DOTATATE (Table S4). The spleen-to-brain and kidney-to-brain mass ratios were in the same range for treated mice as for untreated controls.

**Table S3** Organ masses of mice measured on Day 56 after treatment of mice with 20 MBq [$^{161}$Tb]Tb-DOTA-LM3 or 20 MBq [$^{161}$Tb]Tb-DOTATATE.

| **Treatment** | **Brain** | **Spleen** | **Kidneys** | **Liver** |
|---|---|---|---|---|
| | [mg] | [mg] | [mg] | [mg] |
| Control | 470 ± 20 | 94 ± 6 | 281 ± 19 | 1047 ± 81 |
| [$^{161}$Tb]Tb-DOTA-LM3 | 442 ± 42 | 99 ± 12 | 275 ± 18 | 1161 ± 95 |
| [$^{161}$Tb]Tb-DOTATATE | 420 ± 5 | 96 ± 12 | 279 ± 19 | 1147 ± 110 |

No values of treated mice were significantly different from that of the control group ($p$>0.05).

**Table S4** Organ-to-brain mass ratios determined on Day 56 after treatment of mice with 20 MBq [$^{161}$Tb]Tb-DOTA-LM3 or 20 MBq [$^{161}$Tb]Tb-DOTATATE.

| Treatment | Spleen-to-Brain | Kidneys-to-Brain | Liver-to-Brain |
|---|---|---|---|
| Control | 0.20 ± 0.01 | 0.63 ± 0.03 | 2.2 ± 0.1 |
| [$^{161}$Tb]Tb-DOTA-LM3 | 0.23 ± 0.03 | 0.63 ± 0.05 | 2.6 ± 0.2 |
| [$^{161}$Tb]Tb-DOTATATE | 0.23 ± 0.02 | 0.67 ± 0.07 | 2.8 ± 0.4* |

*Value significantly differed from that of the control group ($p$<0.05).

#### 6.2. Hematological changes

***Purpose:*** Blood cell counts were determined on Day 10, Day 28 and at study end on Day 56 when the mice were sacrificed in order to investigate whether the treatment of mice was well tolerated.

***Methods:*** The methods are described in the main article.

***Results:*** The results are reported mainly in the main article and the determined blood cell counts listed in Tables S5 and S6.

**Table S5** Thrombocyte and red blood cell counts as well as hemoglobin assessed on Day 10, Day 28 and Day 56 after treatment of mice with 20 MBq [$^{161}$Tb]Tb-DOTA-LM3 or 20 MBq [$^{161}$Tb]Tb-DOTATATE.

| Treatment | Thrombocytes | Erythrocytes | Hemoglobin |
|---|---|---|---|
| | [$10^9$ cells/L] | [$10^{12}$ cells/L] | [g/dL] |
| | **Day 10** | | |
| Control | 469 ± 25 | 10.0 ± 0.4 | 13.3 ± 0.6 |
| [$^{161}$Tb]Tb-DOTA-LM3 | 393 ± 58* | 9.6 ± 0.4 | 12.7 ± 0.3 |
| [$^{161}$Tb]Tb-DOTATATE | 403 ± 92 | 9.9 ± 0.3 | 13.0 ± 0.8 |
| | **Day 28** | | |
| Control | 433 ± 109 | 10.1 ± 0.4 | 13.7 ± 0.9 |
| [$^{161}$Tb]Tb-DOTA-LM3 | 465 ± 74 | 9.7 ± 0.3 | 13.1 ± 0.4 |
| [$^{161}$Tb]Tb-DOTATATE | 484 ± 70 | 9.7 ± 0.3 | 12.8 ± 0.5* |
| | **Day 56** | | |
| Control | 499 ± 143 | 10.2 ± 0.2 | 13.2 ± 0.3 |
| [$^{161}$Tb]Tb-DOTA-LM3 | 508 ± 64 | 10.0 ± 0.4 | 13.1 ± 0.7 |
| [$^{161}$Tb]Tb-DOTATATE | 520 ± 68 | 10.1 ± 0.3 | 13.0 ± 0.4 |

*Value significantly differed from that of the control group at the same timepoint ($p$<0.05).

**Table S6** White blood cell counts assessed on Day 10, Day 28 and Day 56 after treatment of mice with 20 MBq [$^{161}$Tb]Tb-DOTA-LM3 or 20 MBq [$^{161}$Tb]Tb-DOTATATE.

| **Treatment** | **Leukocytes** | **Lymphocytes** | **Neutrophils** |
|---|---|---|---|
| | [$10^9$ cells/L] | [$10^9$ cells/L] | [$10^9$ cells/L] |
| | | **Day 10** | |
| Control | 9.2 ± 2.9 | 8.5 ± 2.7 | 0.51 ± 0.32 |
| [$^{161}$Tb]Tb-DOTA-LM3 | 7.7 ± 2.6 | 7.3 ± 2.4 | 0.34 ± 0.22 |
| [$^{161}$Tb]Tb-DOTATATE | 8.1 ± 2.1 | 7.5 ± 2.1 | 0.32 ± 0.14 |
| | | **Day 28** | |
| Control | 10.7 ± 1.3 | 9.9 ± 1.3 | 0.65 ± 0.24 |
| [$^{161}$Tb]Tb-DOTA-LM3 | 7.0 ± 1.1* | 6.2 ± 1.2* | 0.62 ± 0.11 |
| [$^{161}$Tb]Tb-DOTATATE | 7.9 ± 1.2* | 7.2 ± 1.0* | 0.51 ± 0.22 |
| | | **Day 56** | |
| Control | 8.7 ± 1.8 | 8.1 ± 1.8 | 0.46 ± 0.25 |
| [$^{161}$Tb]Tb-DOTA-LM3 | 8.5 ± 1.5 | 7.8 ± 1.3 | 0.41 ± 0.27 |
| [$^{161}$Tb]Tb-DOTATATE | 9.6 ± 1.9 | 8.7 ± 1.5 | 0.63 ± 0.33 |

*Value significantly differed from that of the control group at the same timepoint ($p<0.05$).

#### 6.3. Blood plasma chemistry

***Purpose:*** Several blood plasma parameters were measured at study end to assess the mice for potential signs of liver and kidney damages.

***Methods:*** Blood plasma parameters were determined at study end on Day 56 using blood plasma obtained from centrifuged retrobulbar blood samples collected in lithium-heparin tubes (Microvette, 200 LH, Sarstedt, Germany). The concentrations of blood urea nitrogen, alkaline phosphatase and albumin were determined using a dry chemistry analyzer (DRI-CHEM 4000i, FUJIFILM, Japan) as previously reported [9].

***Results:*** The results are reported mainly in the main article and listed in Table S7. Blood plasma parameters were only determined on Day 56 since retrobulbar blood sampling was approved only on condition to euthanize mice immediately afterwards.

**Table S7** Blood plasma parameters measured on Day 56 after treatment of mice with 20 MBq [$^{161}$Tb]Tb-DOTA-LM3 or 20 MBq [$^{161}$Tb]Tb-DOTATATE.

| **Treatment** | **Blood urea nitrogen** | **Alkaline phosphatase** | **Albumin** |
|---|---|---|---|
| | [mM] | [U/L] | [g/L] |
| Control | 6.3 ± 3.2 | 115 ± 23 | 22 ± 1 |
| [$^{161}$Tb]Tb-DOTA-LM3 | 5.7 ± 2.4 | 106 ± 12 | 21 ± 3 |
| [$^{161}$Tb]Tb-DOTATATE | 6.3 ± 3.0 | 107 ± 22 | 22 ± 1 |

No values were significantly different from that of the control group ($p$>0.05).

### 7. Study II: Tolerability of $^{161}$Tb- and $^{177}$Lu-based radiopeptides (100 MBq/mouse)

#### 7.1. Body masses, organ masses and respective organ-to-brain mass ratios

***Purpose:*** The body mass of mice was monitored as relevant parameter for the general health status of the mice over the entire study. On Day 56, which was defined as the endpoint, potential changes in the organ masses and organ-to-brain mass ratios of mice that received a treatment with one of the respective radiopeptides were assessed in comparison to untreated control mice.

***Methods:*** The relative body mass as well as the organ masses and organ-to-brain mass ratios of mice injected with 100 MBq radiopeptide were determined as reported for Study I.

***Results:*** The average body mass of mice of each group steadily increased over time, regardless of whether the mice were treated with a radiopeptide or belonged to the untreated control group (Fig. S5a). None of the control mice lost more than 5% of the body mass over the entire time of investigation (Fig. S5b). One mouse treated with 100 MBq [$^{161}$Tb]Tb-DOTA-LM3 lost weight at the beginning of the study resulting in 5–10% reduced body mass 6–17 days after radiopeptide injection relative to the body mass on Day 0. The lowest value for the body mass of this mouse was measured on Day 12, however, it fully recovered and reached a relative body mass of >1.0 by study end (Fig. S5c). None of the mice that received [$^{161}$Tb]Tb-DOTATATE experienced a similar loss in body mass (Fig. S5d). Mice that received $^{177}$Lu-labeled peptides showed the opposite situation. While none of the mice that received [$^{177}$Lu]Lu-DOTA-LM3 lost weight (Fig. S5e), one of the mice that received [$^{177}$Lu]Lu-DOTATATE lost >5% of the initial body mass between Day 10 and Day 22 with the lowest mass measured on Day 16 (12% body mass loss; Fig. S5f). This mouse also fully recovered and reached a relative body mass of >1.0 by study end.

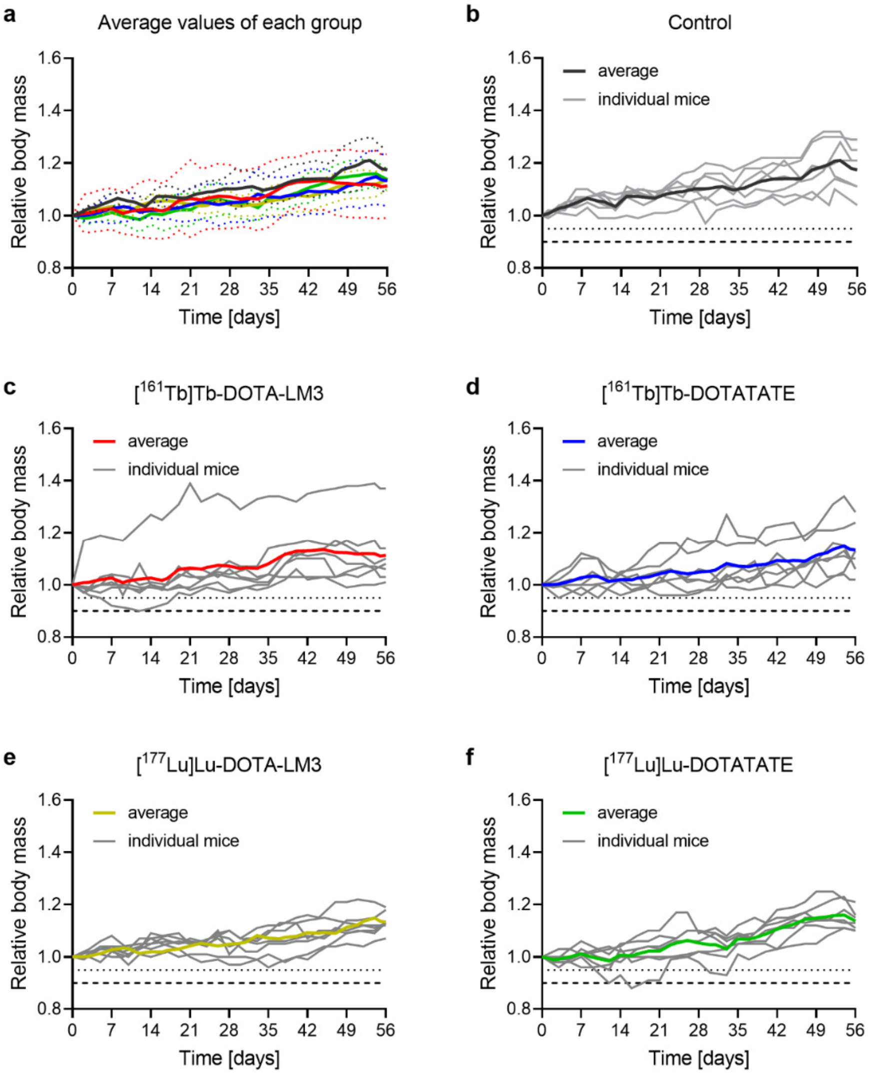


**Fig. S5** Relative body masses of mice injected with 100 MBq (1 nmol) radiopeptide. **a** Relative body masses presented as average ± SD (represented by dotted lines) of the groups in comparison to control mice; **b**−**f** Relative body masses presented for individual mice and as the average presented in color; **b** mice of the untreated control group; **c** mice treated with [$^{161}$Tb]Tb-DOTA-LM3; **d** mice treated with [$^{161}$Tb]Tb-DOTATATE; **e** mice treated with [$^{177}$Lu]Lu-DOTA-LM3; **f** mice treated with [$^{177}$Lu]Lu-DOTATATE. The dotted and dashed lines indicate 5% and 10% body mass loss, respectively, of the initial average body mass of this respective group of mice.

The results of the organ masses and the organ-to-brain mass ratios determined at study end are also described in the main article. The brain masses were highest for control mice but generally equal among mice of all groups. Organ masses for spleen and kidneys were as well in the same range for mice of all groups, however, the liver masses were somewhat higher for mice that received a radiopeptide treatment (1112−1179 mg) than for control mice (1047 ± 81 mg). In the case of mice that received an injection of

[$^{161}$Tb]Tb-DOTATATE, the liver mass was even significantly higher than for control mice (1285 ± 126 mg, *p*<0.05; Table S8). This situation translated in comparable organ-to-brain mass ratios for spleen and kidneys, however, the liver-to-brain mass ratios were higher for treated mice than for untreated controls with significance determined for both $^{161}$Tb-labeled peptides, [$^{161}$Tb]Tb-DOTA-LM3 and [$^{161}$Tb]Tb-DOTATATE (Table S9).

**Table S8** Organ masses of mice measured on Day 56 after application of 100 MBq radiopeptide.

| Treatment | Brain | Spleen | Kidneys | Liver |
|---|---|---|---|---|
| | [mg] | [mg] | [mg] | [mg] |
| Control | 470 ± 20 | 94 ± 6 | 281 ± 19 | 1047 ± 81 |
| [$^{161}$Tb]Tb-DOTA-LM3 | 445 ± 34 | 86 ± 5 | 279 ± 23 | 1179 ± 118 |
| [$^{177}$Lu]Lu-DOTA-LM3 | 436 ± 27 | 81 ± 7 | 269 ± 11 | 1134 ± 77 |
| [$^{161}$Tb]Tb-DOTATATE | 456 ± 17 | 104 ± 11 | 289 ± 20 | 1285 ± 126* |
| [$^{177}$Lu]Lu-DOTATATE | 436 ± 17 | 81 ± 5 | 273 ± 18 | 1112 ± 62 |

*Value significantly differed from that of the control group (*p*<0.05).

**Table S9** Organ-to-brain mass ratios determined on Day 56 after application of 100 MBq radiopeptide.

| Treatment | Spleen-to-Brain | Kidneys-to-Brain | Liver-to-Brain |
|---|---|---|---|
| Control | 0.20 ± 0.01 | 0.63 ± 0.03 | 2.2 ± 0.1 |
| [$^{161}$Tb]Tb-DOTA-LM3 | 0.19 ± 0.02 | 0.63 ± 0.06 | 2.7 ± 0.3* |
| [$^{177}$Lu]Lu-DOTA-LM3 | 0.19 ± 0.01 | 0.62 ± 0.03 | 2.6 ± 0.3 |
| [$^{161}$Tb]Tb-DOTATATE | 0.23 ± 0.03 | 0.64 ± 0.06 | 2.8 ± 0.3* |
| [$^{177}$Lu]Lu-DOTATATE | 0.19 ± 0.01 | 0.63 ± 0.04 | 2.6 ± 0.2 |

*Values significantly differed from that of the control group (*p*<0.05).

#### 7.2. Hematological changes

***Purpose:*** Blood cell counts and blood smears investigated on Day 10, 28 and 56 while bone marrow smears were assessed at study end for each mouse to estimate potential impairment of the bone marrow.

***Methods:*** The methods are described in the main article.

***Results:*** The results are reported below.

#### *7.2.1. Blood cell counts*

The values of blood cell counts are presented in the main article listed in Tables S10 and S11.

**Table S10** Thrombocyte and erythrocyte counts as well as hemoglobin and hematocrit values assessed on Day 10, Day 28 and Day 56 after treatment of mice with 100 MBq [$^{161}$Tb]Tb-DOTA-LM3 or 100 MBq [$^{161}$Tb]Tb-DOTATATE or their respective $^{177}$Lu-labeled counterparts.

| **Treatment** | **Thrombocytes** | **Erythrocytes** | **Hemoglobin** | **Hematocrit** |
|---|---|---|---|---|
| | [$10^9$ cells/L] | [$10^{12}$ cells/L] | [g/dL] | [%] |
| | | **Day 10** | | |
| Control | 469 ± 25 | 10.0 ± 0.4 | 13.3 ± 0.6 | 52 ± 3 |
| [$^{161}$Tb]Tb-DOTA-LM3 | 239 ± 51* | 8.8 ± 0.4* | 12.2 ± 0.7* | 43 ± 4* |
| [$^{177}$Lu]Lu-DOTA-LM3 | 327 ± 60* | 9.4 ± 0.1* | 13.0 ± 0.3 | 45 ± 1* |
| [$^{161}$Tb]Tb-DOTATATE | 387 ± 64 | 9.8 ± 0.4 | 13.4 ± 0.6 | 48 ± 2 |
| [$^{177}$Lu]Lu-DOTATATE | 389 ± 18* | 9.7 ± 0.2 | 13.2 ± 0.3 | 46 ± 2* |
| | | **Day 28** | | |
| Control | 433 ± 109 | 10.1 ± 0.4 | 13.7 ± 0.9 | 50 ± 4 |
| [$^{161}$Tb]Tb-DOTA-LM3 | 448 ± 69 | 9.2 ± 0.1* | 13.0 ± 0.4 | 47 ± 2 |
| [$^{177}$Lu]Lu-DOTA-LM3 | 431 ± 40 | 9.6 ± 0.1 | 13.1 ± 0.2 | 49 ± 2 |
| [$^{161}$Tb]Tb-DOTATATE | 478 ± 106 | 10.2 ± 0.2 | 13.5 ± 0.2 | 49 ± 1 |
| [$^{177}$Lu]Lu-DOTATATE | 483 ± 80 | 10.0 ± 0.3 | 13.5 ± 0.3 | 48 ± 2 |
| | | **Day 56** | | |
| Control | 499 ± 143 | 10.2 ± 0.2 | 13.2 ± 0.3 | 50 ± 4 |
| [$^{161}$Tb]Tb-DOTA-LM3 | 477 ± 74 | 9.6 ± 0.1* | 12.7 ± 0.4 | 47 ± 3 |
| [$^{177}$Lu]Lu-DOTA-LM3 | 463 ± 38 | 9.8 ± 0.2* | 12.9 ± 0.2 | 48 ± 3 |
| [$^{161}$Tb]Tb-DOTATATE | 575 ± 66 | 10.0 ± 0.3 | 12.9 ± 0.7 | 47 ± 2 |
| [$^{177}$Lu]Lu-DOTATATE | 525 ± 47 | 10.2 ± 0.3 | 13.2 ± 0.3 | 49 ± 2 |

*Values significantly differed from that of the control group at the same timepoint ($p<0.05$).

**Table S11** White blood counts assessed on Day 10, Day 28 and Day 56 after treatment of mice with 100 MBq [$^{161}$Tb]Tb-DOTA-LM3 or 100 MBq [$^{161}$Tb]Tb-DOTATATE or their respective $^{177}$Lu-labeled counterparts.

| Treatment | Leukocytes | Lymphocytes | Neutrophils |
|---|---|---|---|
| | [$10^9$ cells/L] | [$10^9$ cells/L] | [$10^9$ cells/L] |
| | | **Day 10** | |
| Control | 9.2 ± 2.9 | 8.5 ± 2.7 | 0.51 ± 0.32 |
| [$^{161}$Tb]Tb-DOTA-LM3 | 4.7 ± 1.1* | 4.1 ± 1.1* | 0.40 ± 0.34 |
| [$^{177}$Lu]Lu-DOTA-LM3 | 5.4 ± 0.6 | 4.9 ± 0.5 | 0.41 ± 0.14 |
| [$^{161}$Tb]Tb-DOTATATE | 5.6 ± 0.7 | 5.2 ± 0.7 | 0.38 ± 0.13 |
| [$^{177}$Lu]Lu-DOTATATE | 5.5 ± 1.0 | 4.9 ± 0.9 | 0.46 ± 0.13 |
| | | **Day 28** | |
| Control | 10.7 ± 1.3 | 9.9 ± 1.3 | 0.65 ± 0.24 |
| [$^{161}$Tb]Tb-DOTA-LM3 | 4.7 ± 1.3* | 4.0 ± 1.2* | 0.58 ± 0.17 |
| [$^{177}$Lu]Lu-DOTA-LM3 | 6.0 ± 1.3* | 5.4 ± 1.1* | 0.40 ± 0.16 |
| [$^{161}$Tb]Tb-DOTATATE | 7.0 ± 1.6* | 6.3 ± 1.6* | 0.52 ± 0.17 |
| [$^{177}$Lu]Lu-DOTATATE | 5.1 ± 1.9* | 4.5 ± 1.5* | 0.45 ± 0.37 |
| | | **Day 56** | |
| Control | 8.7 ± 1.8 | 8.1 ± 1.8 | 0.46 ± 0.25 |
| [$^{161}$Tb]Tb-DOTA-LM3 | 6.0 ± 1.5 | 5.3 ± 1.3* | 0.47 ± 0.23 |
| [$^{177}$Lu]Lu-DOTA-LM3 | 8.5 ± 2.4 | 7.9 ± 2.3 | 0.41 ± 0.18 |
| [$^{161}$Tb]Tb-DOTATATE | 6.2 ± 1.8 | 5.7 ± 1.6 | 0.43 ± 0.31 |
| [$^{177}$Lu]Lu-DOTATATE | 8.1 ± 2.8 | 7.6 ± 2.8 | 0.38 ± 0.19 |

*Values significantly differed from that of the control group at the same timepoint ($p<0.05$).

#### *7.2.2. Blood and bone marrow smears*

The data obtained for blood smears on Day 10, Day 28 and Day 56 are summarized in Table S12. Data with regard to the bone marrow smears are listed in Table S13.

**Table S12** Overview of the assessment of blood smears of mice that received 100 MBq radiopeptide in comparison to findings made for mice of the control group.

| Treatment<br>(Number of mice) | Day 10<br>Observation (Incidence) | Day 28<br>Observation (Incidence) | Day 56<br>Observation (Incidence) |
|---|---|---|---|
| Control (n=7) | None (n=7) | None (n=7) | None (n=7) |
| [$^{161}$Tb]Tb-DOTA-LM3 (n=7) | Less nucleated cells (n=7) | Few neutrophilic myelocyte and/or metamyelocytes (n=2) | Presence of blasts (n=3)<br>Few neutrophilic myelocyte and/or metamyelocytes (n=1) |
| [$^{177}$Lu]Lu-DOTA-LM3 (n=7) | Less nucleated cells (n=7) | Presence of blasts (n=4)<br>Few neutrophilic myelocyte and/or metamyelocytes (n=2) | Presence of blasts (n=4)<br>Few neutrophilic myelocyte and/or metamyelocytes (n=4) |
| [$^{161}$Tb]Tb-DOTATATE (n=7) | Less nucleated cells (n=7) | Presence of blasts (n=1)<br>Few neutrophilic myelocyte and/or metamyelocytes (n=2) | Presence of blasts (n=2)<br>Few neutrophilic myelocyte and/or metamyelocytes (n=2) |
| [$^{177}$Lu]Lu-DOTATATE (n=7) | Less nucleated cells (n=7) | Few neutrophilic myelocyte and/or metamyelocytes (n=1) | Presence of blasts (n=3)<br>Few neutrophilic myelocyte and/or metamyelocytes (n=3) |

**Table S13** Overview of the assessment of bone marrow smears of mice that received 100 MBq radiopeptide in comparison to findings made for mice of the control group.

| Treatment<br>(Number of mice) | Myeloid to erythroid ratio |
|---|---|
| Control (n=7) | 1.11 ± 0.34 |
| [$^{161}$Tb]Tb-DOTA-LM3 (n=7) | 1.74 ± 0.53 |
| [$^{177}$Lu]Lu-DOTA-LM3 (n=7) | 1.23 ± 0.22 |
| [$^{161}$Tb]Tb-DOTATATE (n=7) | 1.83 ± 0.26 |
| [$^{177}$Lu]Lu-DOTATATE (n=7) | 1.98 ± 0.47 |

#### *7.2.3. Differential counts of bone marrow cells*

A differential count of bone marrow cells from selected mice treated with [$^{161}$Tb]Tb-DOTA-LM3 or [$^{161}$Tb]Tb-DOTATATE and control mice was performed. Treated mice had a decreased number of immature precursor cells from the erythrocytic lineage, including prorubricytes, basophilic rubricytes, polychromatophilic rubricytes and metarubricytes. A reduced number of mature lymphocytes was found in bone marrow smears of mice treated with [$^{161}$Tb]Tb-DOTA-LM3, but not in those treated with [$^{161}$Tb]Tb-DOTATATE. On the other hand, treated mice had increased granulocytes, particularly neutrophilic granulocytes. Bone marrow smears from mice treated with [$^{161}$Tb]Tb-DOTA-LM3 showed an increased number of neutrophilic metamyelocytes and neutrophilic band cells while on smears of mice treated with [$^{161}$Tb]Tb-DOTATATE, only neutrophilic band cells were elevated. Additionally, mice treated with [$^{161}$Tb]Tb-DOTA-LM3 or [$^{161}$Tb]Tb-DOTATATE had increased numbers of total monocytic cells. In general, blast cells from the erythrocytic (rubriblasts) and myeloid (myeloblasts and promyelocytes) lineages, as well as neutrophilic myelocytes, prolymphocytes and monoblasts of treated mice were comparable to those of control mice.

### 7.3. Blood plasma chemistry

***Purpose:*** Several blood plasma parameters were measured at study end on Day 56 to assess the mice for potential signs of liver and kidney damages.

***Methods:*** The method was the same as applied for Study I.

***Results:*** The results are reported in the main article and summarized in Table S14. . Blood plasma parameters were only determined on Day 56 since retrobulbar blood sampling was approved only on condition to euthanize mice immediately afterwards.

**Table S14** Blood plasma parameters on Day 56 after treatment with 100 MBq radiopeptide.

| Treatment | Blood urea nitrogen | Alkaline phosphatase | Albumin |
|---|---|---|---|
| | [mM] | [U/L] | [g/L] |
| Control | 6.3 ± 3.2 | 115 ± 23 | 22 ± 1 |
| [$^{161}$Tb]Tb-DOTA-LM3 | 7.8 ± 2.3 | 108 ± 35 | 22 ± 1 |
| [$^{177}$Lu]Lu-DOTA-LM3 | 8.9 ± 0.6 | 95 ± 10 | 20 ± 1 |
| [$^{161}$Tb]Tb-DOTATATE | 8.7 ± 0.8 | 88 ± 21 | 22 ± 3 |
| [$^{177}$Lu]Lu-DOTATATE | 8.0 ± 0.7 | 91 ± 6 | 22 ± 1 |

No values of treated mice were significantly different from those of the control group ($p$>0.05).

#### 7.4. Histological investigations

***Purpose:*** The occurrence of potential early side effects in spleen, kidneys and liver after treatment with 100 MBq $^{161}$Tb- or $^{177}$Lu-labeled somatostatin analogues were assessed based on histo(patho)logical analysis of the respective tissue sections.

***Methods:*** Spleen, kidneys and liver of untreated control mice and mice that received 100 MBq radiopeptide were collected at study end, formalin-fixed and paraffin-embedded. Paraffin blocks were cut into 4-µm thick sections, stained with hematoxylin & eosin (H&E) and assessed by a trained veterinary pathologist using a pre-defined scoring system (Table S15).

**Table S15** Scoring system for histopathological analysis of selected tissues.

| Score | Definition |
|---|---|
| Score 1<br>Minimal | Minor, small or infrequent histopathologic changes ranging from inconspicuous to barely noticeable. For multifocal or diffusely-distributed lesions, this grade was used for processes in which less than approximately 10% of the tissue in an average high-power field was involved. |
| Score 2<br>Slight | Histopathologic changes that are noticeable but not a prominent feature of the tissue. For multifocal or diffusely-distributed lesions, this grade was used for processes in which approximately 10% to 25% of the tissue in an average high-power field was involved. |
| Score 3<br>Moderate | Histopathologic changes that are prominent but not dominant features of the tissue. For multifocal or diffusely-distributed lesions, this grade was used for processes in which approximately 25% to 50% of the tissue in an average high-power field was involved. |
| Score 4<br>Marked | Histopathologic changes that are a dominant but not an overwhelming feature of the tissue. For multifocal or diffusely-distributed lesions, this grade was used for processes in which approximately 50% to 95% of the tissue in an average high-power field was involved. |
| Score 5<br>Severe (Massive) | Histopathologic changes that are an overwhelming feature of the tissue. For multifocal or diffusely-distributed lesions, this grade was used for processes in which more than approximately 95% of the tissue in an average high-power field was involved. |

***Results:*** The results are described in the main article and shown on a representative image which demonstrates the difference between a spleen of an untreated mouse and a spleen of a treated mouse (Fig. S6).

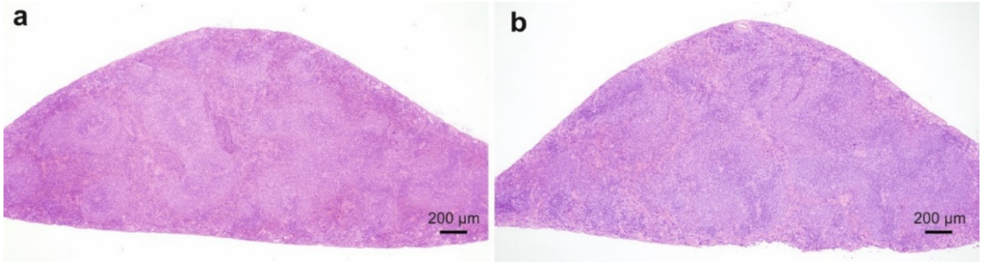


**Fig. S6** **a** Representative image of the spleen of an untreated control mouse; **b** Representative image of the spleen of a treated mouse with lymphoid hyperplasia (visible by the more bluish color).

The lymphoid hyperplasia and subcapsular dilation in the spleen of mice of each group were assessed and scored (Fig. S7a/b). Karyomegaly and tubular basophilia in the kidneys (Fig. S7c/d) as well as karyomegaly in the liver (Fig. S7e) were assessed and scored for mice of each group.

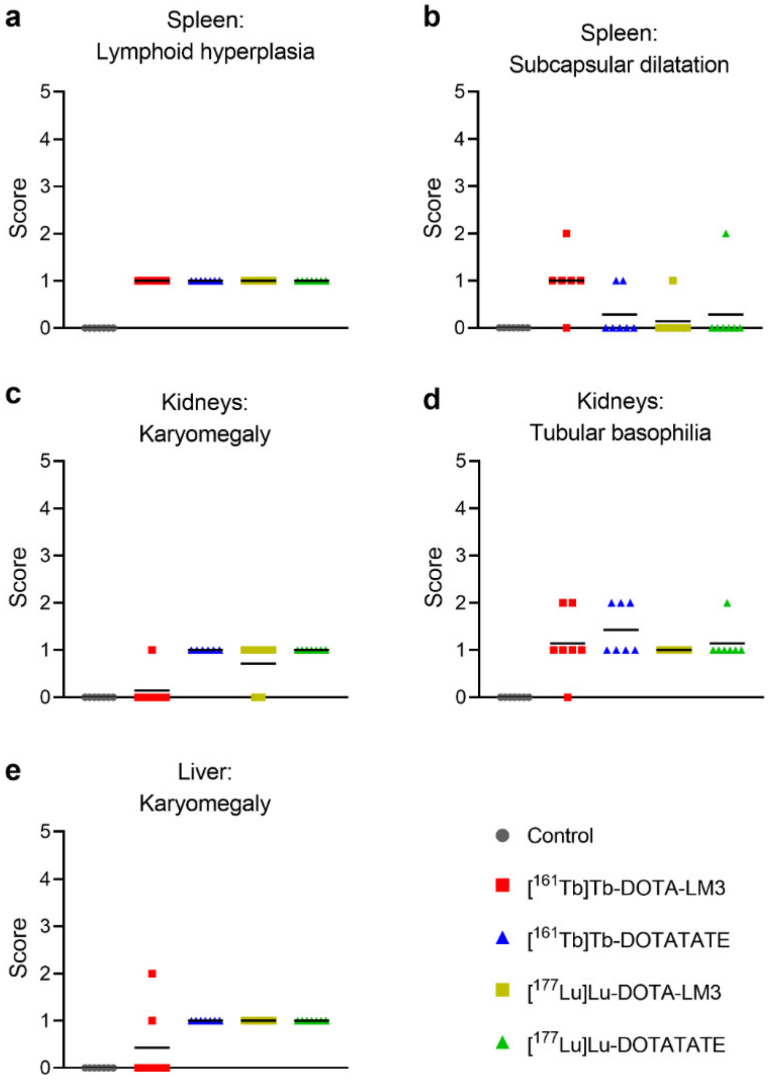


**Fig. S7 a-e** Summary of the most relevant histopathological findings in the spleen, kidneys and liver of mice that received treatment with 100 MBq (1 nmol) radiopeptide in comparison to healthy control mice. **a** Scores assigned to mice with lymphoid hyperplasia; **b** Scores assigned to mice with subcapsular dilatation of blood vessels in the spleen; **c/d** Scores assigned to mice with karyomegaly or tubular basophilia observed in the kidneys; **e** Scores assigned to mice with karyomegaly in the liver.
